# Loss-of-function variants in the schizophrenia risk gene *SETD1A* alter neuronal network activity in human neurons through cAMP/PKA pathway

**DOI:** 10.1101/2021.05.25.445613

**Authors:** Shan Wang, Jon-Ruben van Rhijn, Ibrahim Akkouh, Naoki Kogo, Nadine Maas, Anna Bleeck, Irene Santisteban Ortiz, Elly Lewerissa, Ka Man Wu, Chantal Schoenmaker, Srdjan Djurovic, Hans van Bokhoven, Tjitske Kleefstra, Nael Nadif Kasri, Dirk Schubert

## Abstract

Heterozygous loss-of-function (LoF) mutations in *SETD1A*, which encodes a subunit of histone H3 lysine 4 methyltransferase, were shown to cause a novel neurodevelopmental syndrome and increase the risk for schizophrenia. We generated excitatory/inhibitory neuronal networks from human induced pluripotent stem cells with a *SETD1A* heterozygous LoF mutation (*SETD1A*^+/-^) using CRISPR/Cas9. Our data show that *SETD1A* haploinsufficiency resulted in morphologically increased dendritic complexity and functionally increased bursting activity. This network phenotype was primarily driven by *SETD1A* haploinsufficiency in glutamatergic neurons. In accordance with the functional changes, transcriptomic profiling revealed perturbations in gene sets associated with glutamatergic synaptic function. At the molecular level, we identified specific changes in the cAMP/PKA pathway pointing toward a hyperactive cAMP pathway in *SETD1A*^+/-^ neurons. Finally, by pharmacologically targeting the cAMP pathway we were able to rescue the network deficits in *SETD1A*^+/-^ cultures. Our results demonstrate a link between SETD1A and the cAMP-dependent pathway in human neurons.

## Introduction

Schizophrenia (SCZ) is a complex and heterogeneous syndrome with poorly defined neurobiology. It is a highly heritable disease (~80% heritability) with a substantial genetic component (Hilker et al., 2018; Legge et al., 2021a; Skene et al., 2018). In the past decade, considerable progress has been made to better understand the genetic burden related to SCZ. Genetic loci related to SCZ can either be common variants, which typically exert small effects, or rare variants, which can result in a large effect on individual risk (Legge et al., 2021b). One of the genes with such rare large-effect size variants is *SETD1A*, encoding SET domain-containing protein 1A. Accumulating studies demonstrate that loss-of-function (LoF) and missense mutations in *SETD1A* are associated with SCZ (Singh et al., 2016, 2020; Takata et al., 2014), but also found in individuals with disrupted speech development (Eising et al., 2019) and early-onset epilepsy (Yu et al., 2019). Furthermore, we presented a series of de novo *SETD1A* heterozygous LoF mutations and defined a novel neurodevelopmental syndrome based on a cohort of 15 individuals (age ranging from 34 months to 23 years old) (Kummeling et al., 2020). The core characteristics of these individuals include global developmental delay (such as speech delay or motor delay) and/or intellectual disability, facial dysmorphisms as well as behavior and psychiatric abnormalities, including psychotic episodes. In addition, abnormalities in brain structure, visual and hearing impairments have also been reported in some of these individuals (Kummeling et al., 2020). *SETD1A* mutations thus appear to cause biological vulnerability to a broad neurodevelopmental phenotypic spectrum.

*SETD1A* encodes a subunit of the human Set/COMPASS complex, which methylates histone H3 at position lysine-4 (H3K4me1, H3K4me2, H3K4me3) and participates in the regulation of gene expression. Mouse models with heterozygous LoF mutation of *Setd1a* (*Setd1a^+/-^*) recapitulate SCZ-related behavioral abnormalities, such as deficits in working memory and social interaction (Mukai et al., 2019; Nagahama et al., 2020). At the cellular level, *Setd1a^+/-^* mice display reduced axon branches and dendritic spines, increased neuronal excitability (Mukai et al., 2019) as well as impaired excitatory synaptic neurotransmission (Nagahama et al., 2020). Furthermore, *Setd1a^+/-^* mice show an altered transcriptomic profile in the medial prefrontal cortex (mPFC), a highly relevant region for SCZ (Mukai et al., 2019). In other brain regions, such as visual cortex (V1), *Setd1a^+/-^* mice exhibit aberrant ensemble activity and gamma oscillations (Hamm et al., 2020). All of these studies suggest *SETD1A* haploinsufficiency results in neuronal circuit dysfunction. However, the exact cellular and molecular mechanisms of how *SETD1A* mutations lead to disrupted neuronal connectivity causing such severe mental symptoms, especially in a human context, remain poorly understood.

To investigate the role of SETD1A in neuronal network development and synaptic organization, we generated an isogenic human induced pluripotent stem cell (hiPSC) line with *SETD1A* haploinsufficiency through CRISPR/Cas9. Subsequently, hiPSCs were differentiated into homogenous populations of glutamatergic and GABAergic neurons (Mossink et al., 2021). In *in vitro* cultures containing defined compositions of glutamatergic and GABAergic neurons, we comprehensively analyzed molecular, structural and functional neuronal properties from single cells and neuronal networks during development. The results presented here demonstrate that *SETD1A* haploinsufficiency leads to key morphological, electrophysiological and transcriptional alterations. At molecular level, we show that the *SETD1A^+/-^* network phenotype is mediated by upregulated cyclic adenosine monophosphate (cAMP)/PKA pathway. This has been further confirmed by showing that pharmacological inhibition of cAMP/PKA rescues the *SETD1A^+/-^* network phenotype. Therefore, our results reveal cAMP/PKA as a potential downstream pathway affected by *SETD1A* mutation, opening novel therapeutics opportunities for patients carrying *SETD1A* variant.

## Results

### *SETD1A^+/-^* neuronal networks exhibit dysregulated functional organization

We used CRISPR/Cas9 to generate an isogenic hiPSC line with a heterozygous LoF mutation of *SETD1A* by targeting exon 7 of *SETD1A* in a healthy hiPSC line (Miyaoka et al., 2014) **(Figure 1a)**. We introduced a frameshift mutation in exon 7 leading to a LoF of the protein, mimicking a mutation reported in an individual diagnosed with SCZ (Takata et al., 2014). Two clones with indels in *SETD1A* were selected for further characterization, which carry 28 (clone 1) and 8 (clone 2) base pairs (bp) deletions on one allele, respectively **(Figure 1a, S1a)**, both predicting a premature stop codon. SETD1A mRNA and protein levels were approximately halved in both *SETD1A^+/-^* hiPSC clones, indicating that both variants represent LoF alleles **(Figure 1b-c, S1b)**. All selected clones showed positive expression of pluripotency markers (OCT4, TRA-1-81, NANOG and SSEA4) and karyotyping and off-target analysis was performed to confirm genetic integrity (**Figure S1c-e**).

**Figure 1.**
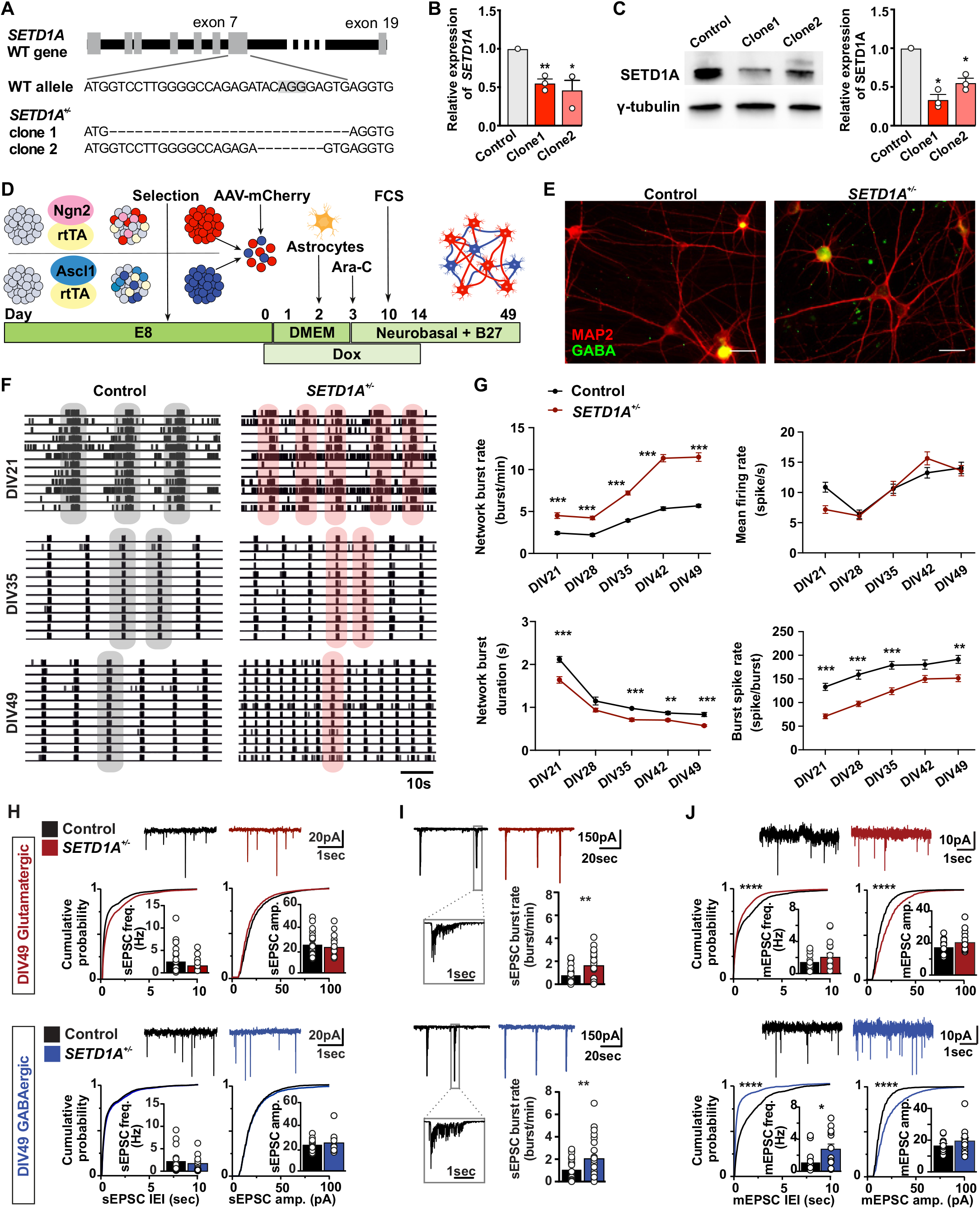
*SETD1A^+/-^* neuronal networks exhibit dysregulated functional organization. (**a**) Generation of *SETD1A* isogenic lines (*SETD1A^+/-^*) using CRISPR/Cas9. Schematic diagram showing the position of sgRNA sequence and indels generated in *SETD1A^+/-^* clone1 and *SETD1A^+/-^* clone2. (**b-c**) qPCR and Western Blot show reduced mRNA (b) and protein level of SETD1A (c). (**d**) Schematic representation of neuronal differentiation workflow. (**e**) Immunofluorescence staining of GABA (green) and MAP2 (red) at DIV49. Scale bar: 30 μm. (**f-g**) Analysis of neuronal activity using MEA recording. (**f**) Representative raster plots (1 min) of electrophysiological activity exhibited by control and *SETD1A^+/-^* neuronal networks at different time points during development. (**g**) Quantification of network parameters as indicated. Sample size: control n = 40 MEA wells. *SETD1A^+/-^* n = 63 MEA wells from 5 independent batches. Representative whole-cell voltage-clamp recordings and quantitative analyses of (**h**) spontaneous postsynaptic currents (sEPSCs) and (**i**) correlated synaptic inputs (sEPSC bursts) in glutamatergic and GABAergic neurons in control and *SETD1A^+/-^* E/I cultures at DIV49. Glutamatergic neurons: control (n = 15), *SETD1A^+/-^* (n = 13); GABAergic neurons: control (n = 15), *SETD1A^+/-^* (n = 12). (**j**) Representative whole-cell voltage-clamp recordings and quantitative analyses of mEPSC activity in glutamatergic and GABAergic neurons. Glutamatergic neurons: control (n = 17), *SETD1A^+/-^* (n = 16); GABAergic neurons: control (n = 16), *SETD1A^+/-^* (n = 13). Data represent means ± SEM. *P < 0.05, **P < 0.01, ***P < 0.001, Students’ T-test with Bonferroni correction for multiple testing (g, h-j) or Kolmogorov-Smirnov test (h) for comparing control vs. *SETD1A^+/-^* cultures.

Disrupted neuronal connectivity has been reported in SCZ patients and animal models of SCZ (Owen et al., 2016). We evaluated whether LoF of *SETD1A* gene results in neuronal network impairment *in vitro*. To this end, we generated composite networks of glutamatergic (~75%) and GABAergic (~25%) neurons (excitatory/inhibitory cultures, E/I cultures) comprised of either control or *SETD1A^+/-^* hiPSCs, by forced expression of the transcription factor *Ngn2* or *Ascl1*, respectively, as recently described (**Figure 1d**) (Mossink et al., 2021). We did not detect any significant differences in the percentage of glutamatergic or GABAergic neurons between control and *SETD1A^+/-^* networks at days *in vitro* (DIV) 49 (**Figure 1e, S2a**). Moreover, following acute treatment with 100 μM Picrotoxin (PTX) at DIV49 on micro-electrode array (MEA), both mean firing rate and network burst duration went up for both control and *SETD1A^+/-^* networks (**Figure S2b-c**), indicating that at DIV49 GABAergic neurons exhibit robust inhibitory control (Mossink et al., 2021).

In order to assess if neuronal network activity differs between control and *SETD1A^+/-^* E/I cultures during development, we used MEA recordings (Frega et al., 2019) (**Figure S2b**). We recorded neuronal network activity once a week from DIV21 to DIV49. After 3 weeks of differentiation, both control and *SETD1A^+/-^* networks showed network burst activity (**Figure 1f**), which indicates the neurons are functionally connected and integrated into a network. Network burst activity increased during development for both groups, and plateaued at around DIV42. Strikingly, *SETD1A^+/-^* networks showed a significantly increased network burst activity compared to control from DIV21 to DIV49, accompanied by a shorter network inter burst interval, a decrease in network burst duration as well as lower spike rate within a burst (**Figure 1g, S2d**). Interestingly, the global activity (i.e., mean firing rate, **Figure 1g**) was similar between control and *SETD1A^+/-^* networks, which implies there was a functional re-organization of network connectivity in *SETD1A^+/-^* E/I cultures, rather than general hyperactivity. Calcium imaging further confirmed the increased synchronized activity in *SETD1A^+/-^* networks (**Figure S3a-d, Supplementary Video1-2**).

We next evaluated if this functional network re-organization in *SETD1A^+/-^* E/I cultures is related to changes in intrinsic properties and/or synaptic inputs using single-cell patch clamp. Intrinsic properties were similar between control and *SETD1A^+/-^* neurons at both DIV21 and DIV49 (**Figure S4, Table S1**), in glutamatergic and GABAergic neurons. We additionally measured general synaptic integration by recording spontaneous excitatory postsynaptic currents (sEPSCs). Neither sEPSC frequency nor amplitude were different between control and *SETD1A^+/-^* cultures at DIV21 or DIV49 (**Figure 1h, Table S1**). However, we did observe that the frequency of temporally correlated bursts of synaptic inputs (sEPSC bursts) onto the postsynaptic neuron was significantly increased at DIV49 in *SETD1A^+/-^* E/I cultures, both on glutamatergic and GABAergic neurons (**Figure 1i**). This is in line with the increased network burst rate at the population level shown in the MEA recordings and confirms that reduced SETD1A expression results in increased network activity. Since analysis of sEPSCs reflects network activity rather than yield quantitative information regarding synaptic connectivity, we next measured miniature postsynaptic currents (mEPSCs), which are action-potential independent and can be indicative for the reorganization of synaptic inputs, altered synapse numbers, and changes in release probabilities (mEPSC frequency) and receptor abundance (mEPSC amplitude). Both mEPSC frequency and amplitude were significantly increased in both glutamatergic and GABAergic *SETD1A^+/-^* neurons at DIV49 (**Figure 1j**). This suggests that networks comprised of *SETD1A^+/-^* neurons not only exhibit increased network activity but also elevated synaptic connectivity, and this could be a major contributor to the network phenotype we observed at the population level.

### *SETD1A^+/-^* neurons show aberrant somatodendritic morphology

Based on the MEA results, we next sought to investigate whether *SETD1A^+/-^* neurons exhibit changes in dendritic morphology and/or synapse formation. At DIV21, control and *SETD1A^+/-^* E/I cultures did not differ from each other in any of the morphological parameters for both glutamatergic and GABAergic neurons (**Figure 2a-b, e-f**). However, at DIV49, *SETD1A^+/-^* neurons displayed a significantly larger soma size, accompanied by a longer dendritic length, more dendritic branches and a larger covered area for both glutamatergic and GABAergic neurons (**Figure 2c-f**). Finally, we performed a Sholl analysis and confirmed that there were no significant differences in the distribution of dendritic length at DIV21 for both neuronal types (**Figure 2a-b**), whereas at DIV49, the dendritic length at multiple distances from the soma was significantly longer in both glutamatergic and GABAergic *SETD1A^+/-^* neurons (**Figure 2c-d**). These results collectively indicate that *SETD1A* haploinsufficiency leads to a more complex somatodendritic morphology, but this only becomes significant during later developmental stages.

**Figure 2.**
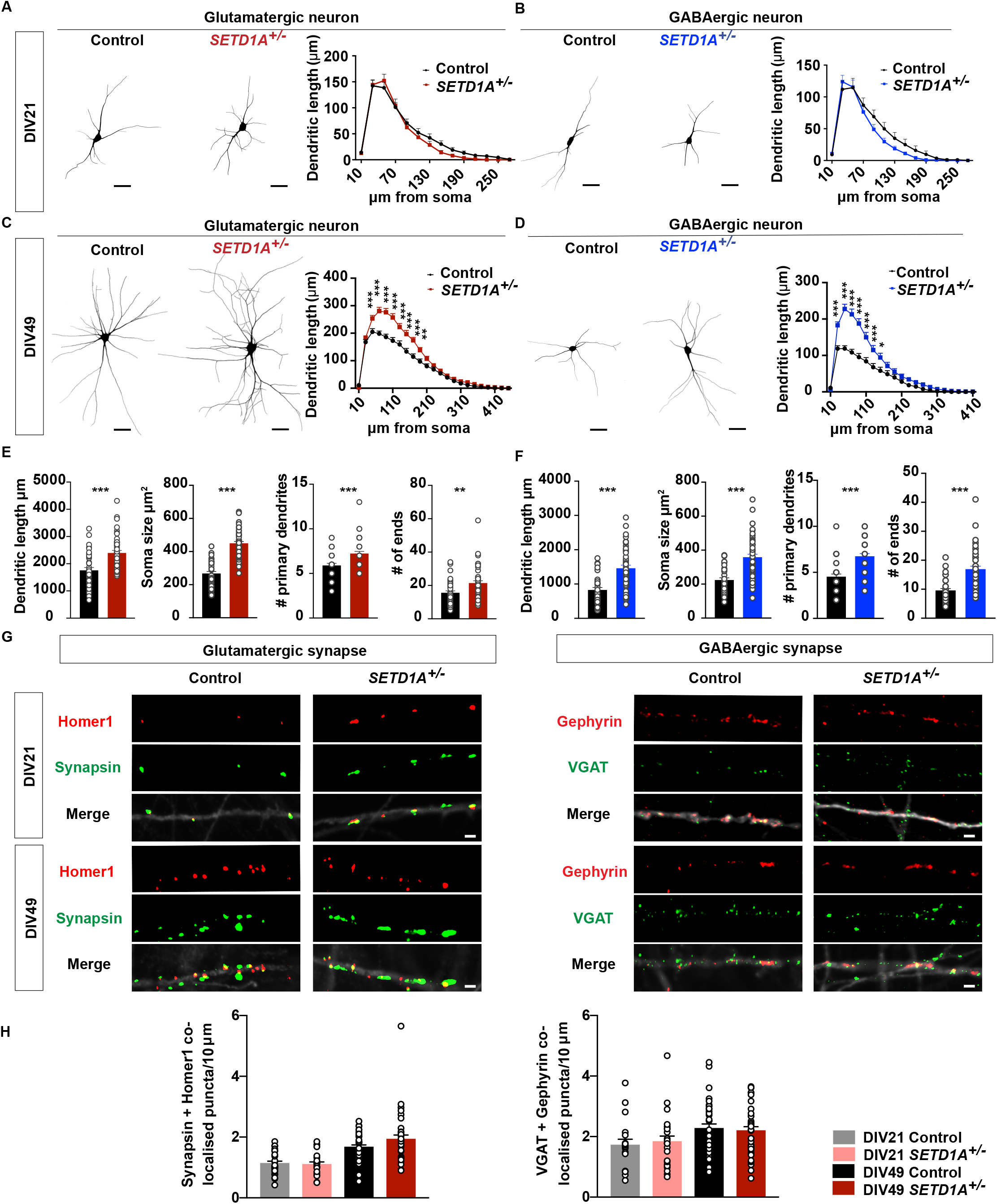
*SETD1A^+/-^* neurons show aberrant somatodendritic morphology. (**a-b**) Representative somatodendritic reconstructions of glutamatergic (a) or GABAergic (b) neurons and Sholl analysis in control and *SETD1A^+/-^* neurons at DIV21 (Glutamatergic neurons: n = 20 for control, n = 20 for *SETD1A^+/-^*; GABAergic neurons: n = 18 for control, n = 21 for *SETD1A^+/-^*). (**c-d**) Representative somatodendritic reconstructions of glutamatergic (c) or GABAergic (d) neurons and Sholl analysis in control and *SETD1A^+/-^* networks at DIV49 (Glutamatergic neurons: n = 38 for control, n = 47 for *SETD1A^+/-^*; GABAergic neurons: n = 33 for control, n = 45 for *SETD1A^+/-^*). (**e-f**) Main morphological parameters in reconstruction for glutamatergic neurons (e) and GABAergic neurons (f) at both DIV21 and DIV49. (**g**) Representative images of immunocytochemistry stained for glutamatergic synapse: Synapsin as a presynaptic marker and Homer1 as a postsynaptic marker, and GABAergic synapse: VGAT as a pre-synaptic marker and Gephyrin as a post synaptic marker. Scale bar = 2 μm (**h**) Quantification of the density of co-localised Synapsin/Homer1 and VGAT/Gephyrin puncta (number per 10 μm). (Synapsin/Homer1: n = 25 for control, n = 21 for *SETD1A^+/-^* at DIV21, n = 34 for control, n = 36 for *SETD1A^+/-^* at DIV49; VGAT/Gephyrin: n = 18 for control, n = 21 for *SETD1A^+/-^* at DIV21, n = 34 for control, n = 36 for *SETD1A^+/-^* at DIV49.) Data represent means ± SEM. *p < 0.05, **p < 0.01, ***p < 0.001, two-way ANOVA with post hoc Bonferroni correction (a-d), unpaired Student’s t test (e-f) or one-way ANOVA with post hoc Bonferroni correction (h).

To measure synapse formation, we immunostained neurons for pre- and postsynaptic markers (Synapsin and Homer1 for glutamatergic synapses, VGAT and Gephyrin for GABAergic synapses). We found no differences in the density of Synapsin/Homer1 or VGAT/Gephyrin co-localised puncta between genotypes at both DIV21 and DIV49 on excitatory and inhibitory neurons (**Figure 2g-h, S5**). These data, combined with the observation that all *SETD1A^+/-^* neurons show a longer dendritic length at DIV49, imply that the total amount of synapses per neuron is higher in *SETD1A^+/-^* neurons, which is reflected by the increase in mEPSC frequency at DIV49 (**Figure 1j**).

### *SETD1A* haploinsufficiency leads to an altered transcriptomic profile

To identify the molecular perturbations that underlie the network phenotypes caused by *SETD1A* haploinsufficiency, we conducted RNA-seq at DIV49. Principal component analysis (PCA) of transcriptional profiles exhibited clear clustering of biological replicates per genotype (**Figure 3a**). We quantified the differentially expressed genes (DEGs) and found that 380 genes were downregulated and 539 genes were upregulated in *SETD1A^+/-^* E/I cultures compared with control cultures (**Figure 3b**). Interestingly, when we performed a disease enrichment analysis using the DisGeNET database which consists of more than 10,000 disorders (Piñero et al., 2020), SCZ was identified as the top hit (**Figure 3c, Table S2**). This indicates that *SETD1A* haploinsufficiency in our model captures some key genetic alterations typically linked to SCZ. In addition, we conducted Gene Ontology (GO) analysis to examine which biological functions are over-represented in the DEGs. Significant enrichment in GO terms relevant to “synaptic function”, “morphogenesis”, “ion channels” and “learning and memory” were identified (**Figure 3d, Table S3**). Proteins encoded by these genes interact with each other as indicated in the protein-protein interaction analysis generated through STRING (Szklarczyk et al., 2021) (**Figure 3e**). Interestingly, we noticed that many GO terms and DEGs relevant to glutamatergic neurons were identified, such as “glutamate receptor binding”, “glutamate receptor signaling pathway” and “glutamatergic synapse” (**Figure 3d-f**). This is consistent with the data from *Setd1a^+/-^* mouse models, where it was shown that SETD1A target genes are highly expressed in pyramidal neurons (Mukai et al., 2019). This also suggests that the altered network phenotype caused by *SETD1A* haploinsufficiency might be driven predominantly by perturbations in glutamatergic neurons.

**Figure 3.**
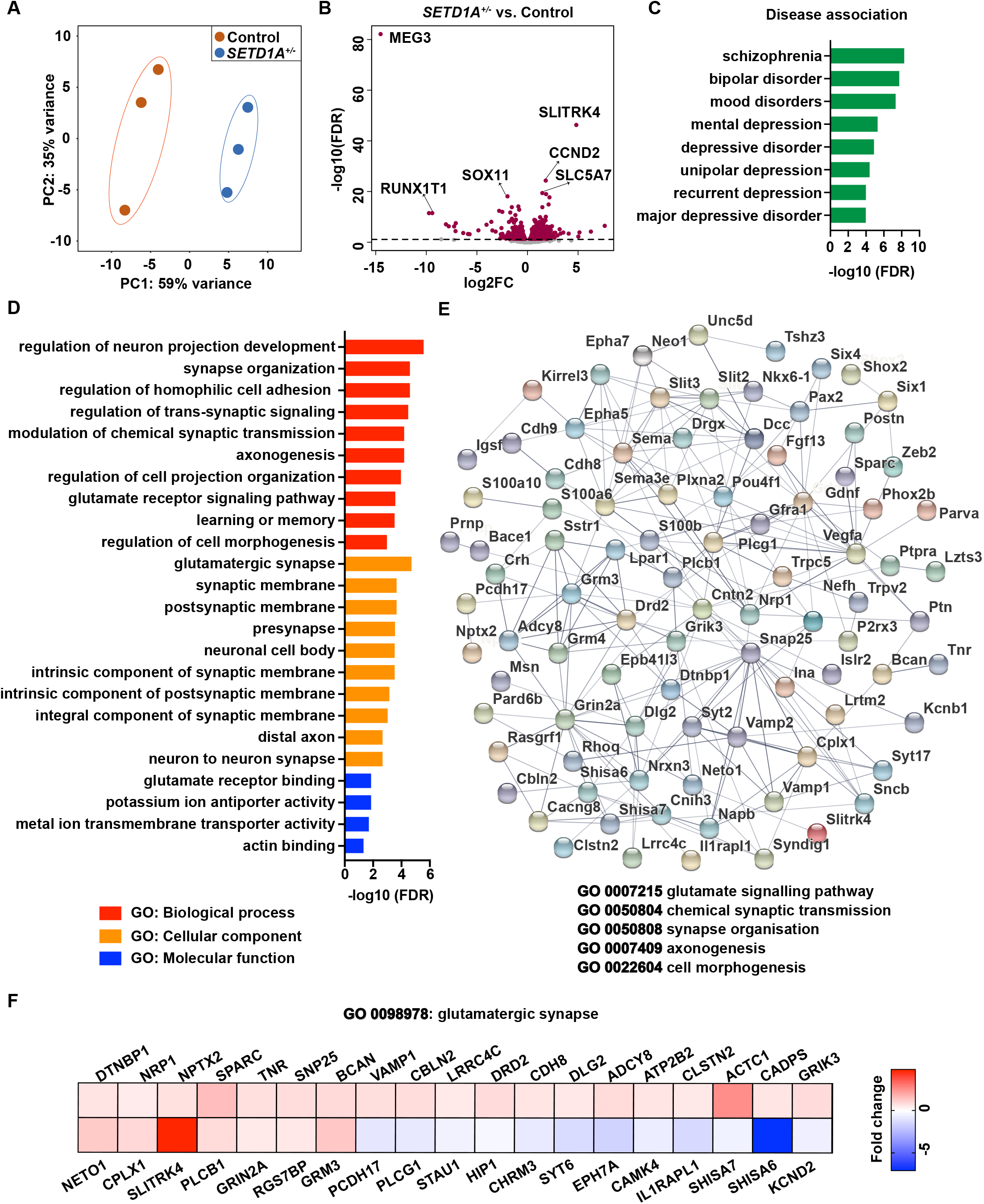
*SETD1A* haploinsufficiency leads to an altered transcriptomic profile. (**a**) PCA showing tight clustering of 3 replicates for each genotype. (**b**) Volcano plots showing differentially expressed genes (DEGs) between *SETD1A^+/-^* and control E/I cultures. Relative to control, significantly up or down-regulated genes are shown in red. Top 3 upregulated and downregulated DEGs are labeled. (**c**) Disease terms of DisGeNET database associated with DEGs. (**d**) Gene Ontology (GO) term analysis of DEGs. (**e**) Diagram for dysregulated protein network showing the interactions among several synaptic functions using STRING. (**f**) Heatmap showing the fold change of DEGs compared to control in gene sets related to glutamatergic synapse.

To explore the cross-species validity of the SETD1A target genes, we compared our DEGs list with the published transcriptome of *Setd1a^+/-^* mouse model from Mukai et al (Mukai et al., 2019), and found in total 11 overlapping genes (**Table S4**). One of these genes is *SLITRK4*, the gene showing the strongest upregulation in *SETD1A^+/-^* cultures (**Figure 3b**), which is involved in neurite outgrowth (Yim et al., 2013). Furthermore, we found a significant association between our DEGs and 78 SFARI genes (Fisher’s exact: p=1.1e^-5^), an autism-related genes database (**Table S5**). This is in line with the suggested genetic overlap between SCZ and autism (Carroll and Owen, 2009). Taken together, these results indicate that transcription is profoundly disturbed in *SETD1A^+/-^* neurons and DEGs are enriched in specific gene sets related to SCZ and synaptic function, especially the ones relevant for glutamatergic synapse function.

### *SETD1A^+/-^* glutamatergic neuronal cultures recapitulate the *SETD1A^+/-^* E/I network phenotype

Our RNA-seq results clearly establish a link between *SETD1A* haploinsufficiency and glutamatergic synapse function. We therefore hypothesized that glutamatergic neurons might be the main contributor to the network phenotype in *SETD1A^+/-^* E/I cultures. To test this hypothesis, we set up homogeneous glutamatergic neuronal cultures (**Figure 4a**). We measured the glutamatergic neuronal network activity on MEA between DIV21 to DIV49 (**Figure 4b**). Consistent with the phenotype of E/I networks, *SETD1A^+/-^* glutamatergic neuronal networks exhibited significantly higher burst activity while decreased burst duration and burst spike rate at DIV49. Global activity (i.e. mean firing rate) did not differ between two groups (**Figure 4c**). Next, we investigated if in glutamatergic cultures, there might be differences between control and *SETD1A^+/-^* neurons at the single cell level (**Figure 4g-k, S8a-b, Table S6**). Similar to the E/I cultures, we observed an increased sEPSC burst rate at the single cell level in particular at DIV49 (**Figure 4j-k, Table S6**). Taken together, our data show that E/I cultures and glutamatergic cultures are similarly affected when SETD1A expression is reduced. This was further supported by the observation that in glutamatergic neuronal networks, *SETD1A^+/-^* neurons showed a similar increase in morphological complexity as in the E/I cultures (**Figure 4d-f, S7a-d**). Finally, we performed a transcriptome analysis for glutamatergic cultures at DIV49. We identified in total 455 DEGs, with 177 downregulated and 278 upregulated genes (**Figure S6a-b**). Notably, we again identified *SLITRK4* as a top hit (**Figure S6b**). Enrichment analysis detected GO terms such as “glutamatergic synapse”, “synapse organization”, “synapse assembly”, and many others related to synaptic function (**Figure S6d-f**). In disease association analysis, we found “bipolar disorder” and “SCZ” within the top hits (**Figure S6c**). Overall, *SETD1A^+/-^* glutamatergic cultures recapitulate the network phenotype observed in the E/I cultures, indicating that SETD1A plays a major role in glutamatergic neurons.

**Figure 4.**
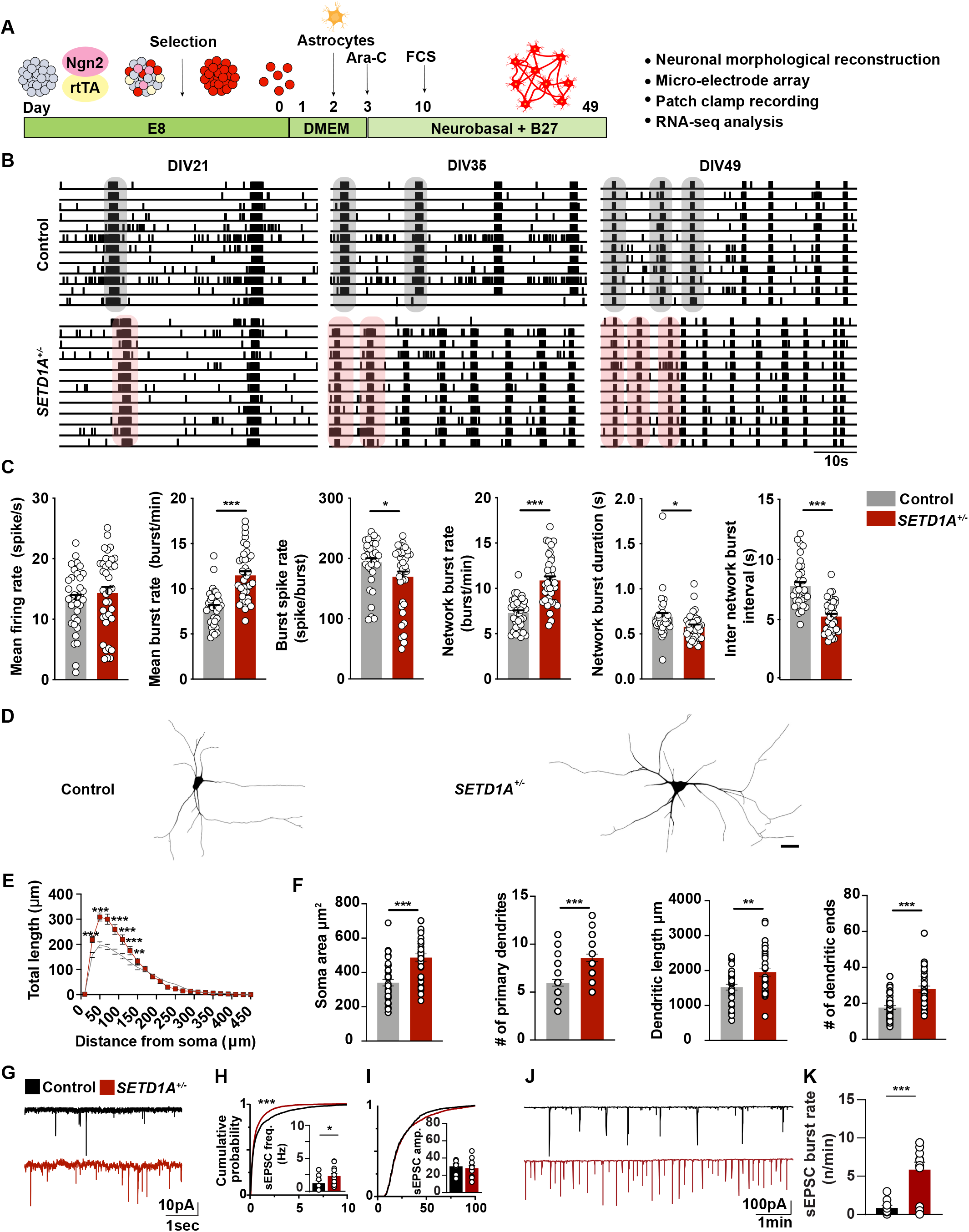
*SETD1A^+/-^* glutamatergic neuronal cultures recapitulate the *SETD1A^+/-^* E/I network phenotype. (**a**) Schematic of neuronal differentiation workflow. (**b**) Representative raster plots (1 min) of electrophysiological activity measured by MEA from control and *SETD1A^+/-^* glutamatergic neuronal networks at different time points during development. (**c**) Quantification of network parameters as indicated at DIV49. Sample size: control n = 35 MEA wells. *SETD1A^+/-^* n = 37 MEA wells from 4 independent batches. (**d-e**) Representative somatodendritic reconstructions of glutamatergic neurons and Sholl analysis in control and *SETD1A^+/-^* networks at DIV 49 (n = 38 for control. n = 33 for *SETD1A^+/-^*). Scale bar = 40 μm. (**f**) Main morphological parameters for reconstructions of glutamatergic neurons at DIV49. (**g**) Representative voltage-clamp recordings of spontaneous inputs (sEPSCs) onto glutamatergic neurons at DIV49 activity and the corresponding quantitative analyses of sEPSC frequency (**h**) and amplitude (**i**). (**j**) Representative voltage-clamp recordings of correlated synaptic inputs (sEPSC bursts) in control and *SETD1A^+/-^* glutamatergic neurons at DIV49 and the corresponding quantitative analyses (**k**). n = 24/17 for control/*SETD1A^+/-^* DIV49. Data represent means ± SEM. *p < 0.05, **p < 0.01, ***p < 0.001, two-way ANOVA with post hoc Bonferroni correction (e), unpaired Student’s t test (c, f) or Students’ T-test with Bonferroni correction for multiple testing (h, i, k).

### *SETD1A* haploinsufficiency leads to activation of cAMP/PKA/CREB pathway

The changes in neuronal network activity observed in *SETD1A^+/-^* cultures (E/I and glutamatergic cultures) suggest that pathways related to regulation of neuronal activity could be affected. Interestingly, in our transcriptome data we found that the upregulated DEGs were enriched in several annotations related to second messenger signaling, such as “G protein-coupled receptor signaling pathway, coupled to cyclic nucleotide second messenger”, “second-messenger-mediated signaling”, and “adenylate cyclase-modulating G protein-coupled receptor signaling pathway” (**Figure 5a, Table S7**). In agreement with this finding, we found that multiple enzymes involved in the synthesis and degradation of cAMP were dysregulated, both in the E/I and glutamatergic cultures. Specifically, genes coding for proteins involved in cAMP production (*ADCY2*, *ADCY3* and *ADCY8*) (Pieroni et al., 1995; Qiu et al., 2016; Wang and Storm, 2003) were significantly upregulated, whereas genes encoding for phosphodiesterases (*PDE12*, *PDE7a* and *PDE1a*), important for cAMP degradation (Epstein, 2017; Evripioti et al., 2019; Keravis and Lugnier, 2012), were downregulated (**Figure 5b**). This strongly suggests that the cAMP pathway is hyperactive in *SETD1A^+/-^* neurons.

**Figure 5.**
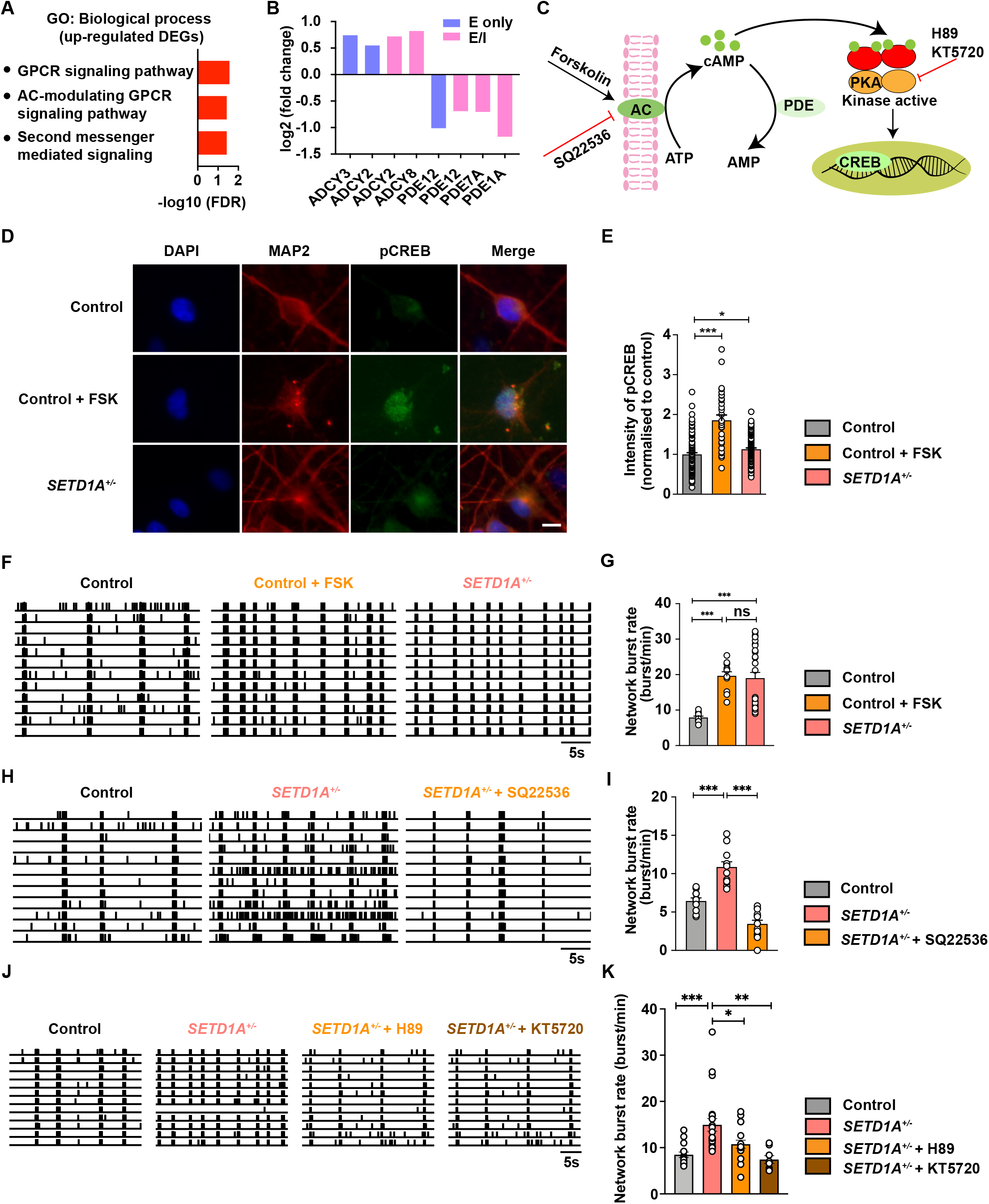
*SETD1A* haploinsufficiency leads to activation of cAMP/PKA/CREB pathway. (**a**) GO terms related to second messenger signaling associated with upregulated differentially expressed genes (DEGs) (**b**) Change of mRNA levels of genes coding for adenylyl cyclase (AC) and phosphodiesterases. (**c**) Schematic of cAMP/PKA/CREB pathway and drug target at different steps. (**d-e**) Representative images showing pCREB expression and quantification of intensity of pCREB for control (n = 105 cells), control + forskolin (n = 30 cells) and *SETD1A^+/-^* neurons (n = 122 cells from 2 independent batches). Scale bar = 10 μm. (**f-g**) Representative raster plot for 30 sec recorded by MEA and quantification of network burst rate showing the effect of AC agonist forskolin on control networks (sample size: control n=10 wells, control + forskolin n = 12 wells, *SETD1A^+/-^* n = 28 wells). (**h-i**) Representative raster plot for 30 sec recorded by MEA and quantification of network burst rate showing the effect of AC inhibitor SQ22536 on *SETD1A^+/-^* E/I cultures (sample size: control n=11 wells, *SETD1A^+/-^* + SQ22536 n = 12 wells, *SETD1A^+/-^* n =13 wells). (**j-k**) Representative raster plot for 30 sec recorded by MEA and quantification of network burst rate showing the effect of PKA inhibitor H89 and KT5720 on *SETD1A^+/-^* E/I cultures (sample size: control n=15 wells, *SETD1A^+/-^* + H89 n = 18 wells, *SETD1A^+/-^* + KT5820 n = 18 wells, *SETD1A^+/-^* n = 22 wells). Data represent means ± SEM. *P <0.05, **P < 0.01, ***P < 0.001, one-way ANOVA test and post-hoc Bonferroni correction.

To test this hypothesis, we measured the phosphorylation of cAMP-response element binding protein (pCREB), which is a downstream signaling protein in the cAMP/PKA pathway (Tasken et al., 1993) (**Figure 5c**). We immunostained neurons from DIV49 E/I cultures for anti-pCREB, an antibody detecting endogenous levels of CREB only when phosphorylated at serine^133^. As a positive control, we stimulated the cAMP pathway on control E/I cultures with 10 μM forskolin (FSK), a well-known adenylyl cyclase (AC) agonist and PKA/CREB pathway activator (Delghandi et al., 2005; Namkoong et al., 2009; Schiller et al., 2010). As expected, FSK significantly increased the level of pCREB in control neurons. In addition, we found that the level of pCREB in *SETD1A^+/-^* neurons was significantly higher compared to control neurons (**Figure 5d-e**). This result supports our hypothesis that the cAMP/PKA/CREB pathway exhibits increased activity in *SETD1A^+/-^* neurons.

To further investigate the link between enhanced cAMP activation and the neuronal network phenotype observed in *SETD1A^+/-^* neurons, we bidirectionally modulated cAMP levels in control or *SETD1A^+/-^* neurons from E/I cultures. Interestingly, we found that acute stimulation of DIV51 control neurons with 10 μM FSK for 1 hour resulted in an increase in network bust rate, with a parallel reduction in inter network burst interval (**Figure 5f-g, S9a-c**), mimicking the network phenotype of *SETD1A^+/-^* E/I cultures. In addition, burst spike rate and network burst duration were decreased, although the difference was not significant (**Figure S9c**). These results show that control networks treated with forskolin resemble *SETD1A^+/-^* neuronal networks. In a second set of experiments, we examined whether blocking cAMP using AC inhibitor SQ22536 in *SETD1A^+/-^* networks (starting from DIV42) would be sufficient to normalize the phenotype caused by *SETD1A* haploinsufficiency to control levels. Indeed, after 8 days of treatment with 100 μM SQ22536, we found that the major network parameters were normalized, including the network burst rate (**Figure 5h-i, S10a-b**). We next investigated whether acute manipulation of the PKA pathway, which is a downstream target of cAMP, would be sufficient to normalize *SETD1A^+/-^* network activity to control levels. Two independent chemical inhibitors of PKA, H89 and KT5720 (Murray, 2008; Song et al., 2015), were applied to *SETD1A^+/-^* networks at DIV51 on MEA. A one-hour treatment with either H89 (2 μM) or KT5720 (1 μM) in *SETD1A^+/-^* networks, was sufficient to normalize the major network parameters, including the network burst rate (**Figure 5j-k, S11a-c**). Taken together, our results suggest that cAMP/PKA is the main molecular pathway through which *SETD1A* haploinsufficiency leads to key neuronal network alteration.

## Discussion

Heterozygous LoF variants in *SETD1A*, with high penetrance for SCZ, provides a unique opportunity to investigate neuronal dysfunction that might underlie the increased vulnerability to SCZ. Here, we reveal crucial molecular signatures and neurodevelopmental abnormalities caused by *SETD1A* haploinsufficiency in human neurons. We show that *SETD1A* haploinsufficiency leads to altered neuronal network organization, within which changes in glutamatergic neuronal function might be one of the main contributors. Furthermore, we identify increased cAMP/PKA activity as a molecular mechanism to the functional phenotype of *SETD1A^+/-^* networks.

One major consistent characteristic of hiPSC-derived *SETD1A^+/-^* networks is the increased network burst rate, which is reflected by an increase in synaptic connectivity at the single cell level. Synchronized burst activity plays an important role during early brain development (Khazipov and Luhmann, 2006) and is considered to be a fundamental mechanism for information processing, hence relevant for perception, memory and cognition (Lisman, 1997). Intriguingly, abnormalities in neural synchronization *in vivo*, which reflect neural circuit dysfunction, are suggested as one of the core pathophysiological mechanisms in SCZ (Ford et al., 2007; Uhlhaas and Singer, 2010). For example, increased oscillatory synchronization is found to be correlated with positive symptoms of SCZ (Lee et al., 2006; Spencer et al., 2009), while both increased and reduced oscillations are associated with negative symptoms (Lee et al., 2003; Uhlhaas and Singer, 2010). In the primary visual cortex of *Setd1a^+/-^* mice, Hamm et al identified reduced oscillations, providing circuit-level insight into the underlying neurobiology of sensory-processing abnormality seen in SCZ (Hamm et al., 2020). In this regard, both our human *in vitro* model and mouse model showed aberrant synchronized activity at different developmental stages, suggesting circuit disruption caused by *SETD1A* haploinsufficiency.

Disrupted glutamatergic signaling plays a critical role in the pathogenesis of SCZ (Tsai and Coyle, 2002). Consistently, on several levels, our data provide evidence supporting the idea that glutamatergic signaling are one of the main contributors for the network phenotype caused by *SETD1A* haploinsufficiency. First, through transcriptional profiling of *SETD1A^+/-^* E/I cultures, we identified a genetic signature suggestive of perturbed function of glutamatergic signaling. Dysregulated genes include the ones encoding for metabotropic glutamate receptors (*GRM4*, *GRM3*), ionotropic glutamate receptors subunits (*GRIK3*, *GRIN2A*), as well as proteins involved in glutamatergic signaling (*SHISA6*, *SHISA7*, *CAMK4*, *PSD93*, *VAMP1*). This is in line with the previous finding that target genes of SETD1A were mainly expressed in pyramidal neurons (Mukai et al., 2019). In particular, *SHISA6* which is important for AMPA receptor function has been strongly down-regulated, suggesting dysregulated AMPA receptor activity (Klaassen et al., 2016). Furthermore, in L2/3 pyramidal neurons from the mPFC of *Setd1a^+/-^* mice, only excitatory synaptic transmission was altered, whereas inhibitory synaptic transmission remained unchanged (Nagahama et al., 2020). Second, in E/I networks, when GABAergic inputs were blocked by PTX, the phenotype of increased networks burst frequency still remained (**Figure S2c**). In addition, measurements from single-cell electrophysiology and MEA with homogenous glutamatergic neurons show the same network signature as observed in E/I networks, indicating glutamatergic neurons are one of the main driving factors in this phenotype. Third, from both E/I and glutamatergic cultures, transcriptomic data highlight a striking upregulation of *SLITRK4*, which is involved in neurite outgrowth and synaptogenesis, especially in glutamatergic synapse formation (Yim et al., 2013). Moreover, it has been shown that variants in *SLITRK4* are associated with neuropsychiatric disorders, such as schizophrenia (Kang et al., 2016). *SLITRK4* has also been identified as a DEG from *Setd1a^+/-^* mice (Mukai et al., 2019), emphasizing its important role as a downstream target of SETD1A across species. Indeed, our data show that *SETD1A* haploinsufficiency leads to increased dendritic complexity in human glutamatergic neurons. This morphological phenotype however was not detected in the *SETD1A^+/-^* mice model from Mukai et al. where pyramidal neurons in mature prefrontal cortical networks showed unchanged dendritic arborization (Mukai et al., 2019). This difference can potentially be explained by the differences in the assessed developmental stages or species related differences in SETD1A function. Taken together, these results imply that SETD1A plays an essential role in the function of glutamatergic signaling. However, dysfunction of GABAergic signaling might be involved in the network phenotype as well. Our results show that *SETD1A* haploinsufficiency leads to increased dendritic complexity of GABAergic neurons, which likely affects the input connectivity and signal integration of these neurons. Further experiments are needed to clearly dissect the role of SETD1A in GABAergic neurons.

Our data identify several molecular pathways that may contribute to the increased network burst frequency in *SETD1A^+/-^* networks. In particular, our transcriptomic data indicated that *SETD1A* haploinsufficiency leads to increased cAMP/PKA, which has been shown to enhance synaptic strength through different aspects, such as by modulating different ion channels (Anderson et al., 2000; Beaumont and Zucker, 2000; Connors et al., 2008; Gutierrez-Castellanos et al., 2017; Hell’ et al., 1995; van der Horst et al., 2020; Renner et al., 2017) and increasing neurite outgrowth (Aglah et al., 2008; Chu et al., 2006; Wan et al., 2011). Intriguingly, a recent study demonstrated that cAMP/PKA induces calcium influx through voltage-gated calcium channel, which pushes the neuronal network towards a large-scale and synchronized burst activity (Thornquist et al., 2021). Consistently, our calcium imaging data showed increased synchronized activity in *SETD1A^+/-^* networks, suggesting that there could be a similar cAMP-triggered mechanism in our model.

cAMP signaling has been implicated in neurodevelopmental and neuropsychiatric disorders, including SCZ (Funk et al., 2012; Millar et al., 2005; Muly, 2002; Turetsky and Moberg, 2009; Vacic et al., 2011; Wang et al., 2018), bipolar disorder (Chang et al., 2003; Ren et al., 2014), Fragile X syndrome (Berry-Kravis et al., 1995; Berry-Kravis and Huttenlocher, 1992; Kelley et al., 2007) and autism (Kelley et al., 2008;

Zamarbide et al., 2019). For example, in patients with bipolar disorder, cAMP/PKA signaling is upregulated (Chang et al., 2003), while in the case of Fragile X syndrome, cAMP/PKA signaling is decreased (Berry-Kravis et al., 1995). These findings emphasize the essential role of balanced cAMP activity in normal brain function, making it an appealing therapeutic target. Interestingly, our results show that on network level both blocking the PKA pathway and inhibiting AC can rescue the phenotype. However, we cannot rule out the possibilities that other downstream effectors of cAMP such as EPAC (a guanine-nucleotide-exchange factor) and cyclic-nucleotide-gated ion channels (Sassone-Corsi, 2012), are also partly involved in the phenotype caused by *SETD1A* haploinsufficiency. In addition, since cAMP/PKA signaling functions in various cells types, therapeutic intervention of its activity therefore needs further clarification of its cell type specific roles. In this context, future studies should comprehensively investigate whether upregulated cAMP/PKA signaling caused by *SETD1A* haploinsufficiency also applies to other brain regions or other cell types related to SCZ or neurodevelopmental disorders, such as serotonergic (Carvajal-Oliveros and Campusano, 2021) or dopaminergic neurons (Eells, 2005), as well as glial cells (Blanco-Suárez et al., 2017).

Even though our current human neuronal *in vitro* model skips certain early developmental stages, such as the neuro-progenitor cell stage, of neuronal networks and it cannot capture the cellular complexity of fully mature cortical networks, the defined composition of human neuronal networks allows the assessment of core molecular and cellular mechanisms related to *SETD1A* haploinsufficiency. Taken together, our data suggest glutamatergic synaptic dysfunction is one of the potential pathogenic mechanisms of *SETD1A* haploinsufficiency-associated disorders and we identified cAMP/PKA dysregulation as underlying mechanisms responsible for the altered network phenotype in *SETD1A^+/-^* cultures. Future studies, using both rodent models and human *in-vitro* neuronal models are required to further explore the impact and therapeutic potential of cAMP/PKA in *SETD1A*-associated disorders.

## Materials and Methods

### Cell culture and neuronal differentiation

hiPSCs used in this study were obtained from reprogrammed fibroblasts (Miyaoka et al., 2014). hiPSCs were cultured on a 6-well plate pre-coated with 1:15 (diluted in DMEM/F12 medium) Matrigel (Corning, #356237) in Essential 8 Flex medium (Thermo Fisher Scientific) supplemented with primocin (0.1 g/ml, Invitrogen) at 37°C/5%CO_2_. hiPSCs were infected with *Ascl1* or *Ngn2* and *rtTA* lentivirus (The transfer vector used for the rtTA lentivirus is pLVX-EF1α-(Tet-On-Advanced)-IRES-G418(R); The transfer vector used for the Ngn2 lentivirus is pLVX-(TRE-thight)-(MOUSE) Ngn2-PGK-Puromycin(R); The transfer vector used for the Ascl1 lentivirus is pLV[TetOn]-Puro-TRE>mAscl1. All the plasmids are available upon request). Medium was supplemented with puromycin (0.5 g/ml) and G418 (50 g/ml). Medium was refreshed every 2-3 days and hiPSCs were passaged twice per week using an enzyme-free reagent (ReLeSR, Stem Cell Technologies).

The neuronal differentiation protocol used in this article was previously described (Mossink et al., 2021). Glutamatergic neurons were either cultured alone or in coculture with GABAergic neurons. When cocultured, GABAergic neurons were plated at days *in vitro* (DIV) 0 and transduced with AAV2-hSyn-mCherry (UNC Vector Core) for visualization. After 4 hours incubation, cultures were washed twice with DMEM/F12 (Thermo Fisher Scientific) after which glutamatergic neurons were plated into the well. hiPSCs were plated in E8 flex supplemented with doxycycline (4 μg/ml), Revitacell (1:100, Thermo Fisher Scientific) and Forskolin (10 μM, Sigma). At DIV 1 cultures were switched to DMEM/F12 containing Forskolin, N2 (1:100, Thermo Fisher Scientific), non-essential amino acids (1:100, Sigma), primocin (0.1 μg/ml, Invivogen), NT3 (10 ng/ml, PromoCell), BDNF (10 ng/ml, PromoCell), and doxycycline (4 μg/ml). To support neuronal maturation, freshly prepared rat astrocytes were added to the culture in a 1:1 ratio at DIV 2. At DIV 3 the medium was changed to Neurobasal medium (Thermo Fisher Scientific) supplemented with Forskolin (10 μM, Sigma), B-27 (Thermo Fisher Scientific), GlutaMAX (Thermo Fisher Scientific), primocin (0.1 μg/ml), NT3 (10 ng/ml), BDNF (10 ng/ml), and doxycycline (4 μg/ml). To remove the proliferating cells from the culture, cytosine-b-D-arabinofuranoside (Ara-C, 2 μM, Sigma) was added to the medium at DIV 3. From DIV 6 onwards half of the medium was refreshed three times a week. From DIV 10 onwards, the medium was additionally supplemented with 2.5% FBS (Sigma) to support astrocyte viability. After DIV 13, Forskolin and doxycycline were removed from the medium. Cultures were kept at least until DIV49.

### CRISPR/Cas9 editing of *SETD1A*

CRISPR/Cas9 technology was used to create a heterozygous indel mutation in exon 7 of *SETD1A* in a healthy hiPSC line derived from a male, 30 years old (Mandegar et al., 2016). In brief, sgRNAs were designed to specifically target *SETD1A* (GTCCTTGGGGCCAGAGATAC AGG), and then cloned into pSpCas9(BB)-2A-Puro (PX459) V2.0 (Addgene #62988). Single-cell suspension of hiPSCs was nucleofected with 5 μg of the generated SpCas9-sgRNA plasmid using the P3 Primary Cell 4D-Nucleofector Kit (Lonza, #V4XP-3024) in combination with the 4D Nucleofector Unit X (Lonza, #AAF-1002X), program CA-137. After nucleofection, cells were resuspended in E8 Flex supplemented with Revitacell and seeded on Matrigel pre-coated plates. One day after the nucleofection, 0.5 μg/ml puromycin was added for 24 hours for selection. Puromycin-resistant colonies were manually picked and Sanger Sequencing was performed to ensure heterozygous editing of exon 7. Two positive clones were selected for further characterization. Cells were tested routinely as mycoplasma-negative. The expression of pluripotency markers OCT4, NANOG, SSEA4 and TRA1-81 were detected with immunocytochemistry. Karyotyping was performed as a service by the Diagnostics Department at Radboud University Medical Center. Potential off-target sites were checked by sanger sequencing.

### Immunocytochemistry

Cells seeded on coverslips were fixed with 4% paraformaldehyde supplemented with 4% sucrose for 15 min at room temperature, followed by permeabilization with 0.2% triton for 10 min. Nonspecific binding sites were blocked by incubation in 5% normal goat serum for 1 hour at room temperature. Cells were incubated with primary antibodies overnight at 4°C. The second day, secondary antibodies, conjugated to Alexa-fluorochromes, were added and incubated for 1 hour at room temperature. Hoechst 33342 was used to stain the nucleus before cells were mounted with fluorescent mounting medium (DAKO, #S3023). The following primary antibodies were used: Rabbit anti-MAP2 (1:1000, Abcam, #ab32454); Mouse anti-Gephyrin (1:1000, Synaptic Systems 147011); Rabbit anti-VGAT (1:500, Synaptic systems 131013); Guinea pig anti-Synapsin 1/2 (1:1000, Synaptic Systems 106004); Mouse anti-Homer1 (1:500, Synaptic Systems 160011); Rabbit anti-Phospho-CREB (1:500, Cell Signaling 87G3 9198). Secondary antibodies that were used are: Goat-anti rabbit Alexa 568 (1:1000, Invitrogen, A11036); Goat-anti-mouse Alexa 488 (1:1000, Invitrogen, A11029); Goat anti-guinea pig Alexa Fluor 568 (1:2000, Invitrogen, A11075); Goat anti-guinea pig Alexa Fluor 647 (1:1000, Invitrogen, A21450). Cells were imaged at 63x magnification using the Zeiss Axio Imager Z1 equipped with ApoTome. Fluorescent images were analyzed using FIJI software (Schindelin et al., 2012). Synapse puncta were counted manually and normalized to the length of the dendritic branch where they reside.

### Western Blot

To lyse the cells, medium was removed and the well was washed with 2 ml ice cold PBS before 100 μl lysis buffer were applied (RIPA buffer supplemented with PhosSTOP; Roche) and protease inhibitors (complete Mini, EDTA free; Roche). Before blotting the protein, concentration was determined by means of a Pierce^™^ BCA protein assay (Thermo Scientific 23225). For each sample, the same amount of protein around 15 μg was loaded and separated by SDS-PAGE. Depending on the primary antibody, separated proteins were transferred to PVDF membrane (BioRad). Primary antibodies were used are: Mouse anti-SET1A (1: 500, Santa Cruz 515590); Mouse anti-γ-tubulin (1:1000, Sigma T5326). For visualization horseradish peroxidase-conjugated secondary antibodies were used: Goat anti-mouse (1:20000; Jackson Immuno Research Laboratories 115-035-062).

### Neuron reconstruction

Reconstruction was performed using Neurolucida 360 (Version 2017.01.4, Microbrightfield Bioscience). Neurons were fixed and labeled with MAP2 antibody. To distinguish GABAergic neurons from glutamatergic neurons, we infected GABAergic neurons with AAV2-hSyn-mCherry for (UNC Vector Core) visualization. We chose two time points to fix the neurons: DIV21 and DIV49. This allows us to compare the morphological phenotype at different developmental stages. Fluorescent images of MAP2-labelled neurons were taken at 20x magnification using Zeiss Axio Imager Z1 equipped with ApoTome. The images were stitched using Fiji 2018 software with the stitching plugin and followed by reconstruction using Neurolucida 360 (Version 2017.01.4, Microbrightfield Bioscience). The 3-dimensional reconstructions and quantitative morphometrical analyses focused on the somatodendritic organization of the neurons. The axon was excluded based on its long, thin properties and far-reaching projections. Neurons that had at least two primary dendrites were selected for reconstruction and further analysis. For morphometrical analysis, we determined soma size, number of primary dendrites, dendritic nodes and ends and total or mean dendritic length as well as covered surface by dendritic trees. In addition, we also investigated dendritic complexity by performing Sholl analysis. Sholl profile was obtained by applying a series of concentric circles at 20 μm interval from the soma center, subsequently, dendritic length, number of intersections and number of nodes of the neurons were measured for each distance interval.

### MEA recordings and analysis

All recordings were performed using the 24-wells MEA system (Multichannel Systems, MCS GmbH, Reutlingen, Germany). Recordings and analysis were performed according to previous published protocols (Frega et al., 2019). Briefly, spontaneous electrophysiological activity of hiPSC-derived neuronal networks cultured on MEA was recorded for 10 minutes in a recording chamber which was constantly maintained at 37°C with 95%O_2_ and 5%CO_2_. Before recording, cultures on MEA were allowed to adapt for 10 min in the recording chamber. The recording was sampled at 10 kHz, and filtered with a high-pass filter with a 100 Hz cut-off frequency and a low-pass filter with a 3500 Hz cut-off frequency. The spike detection threshold was set at ± 4.5 standard deviations. Spike, burst and network burst detection was performed by a built-in algorithm in Mulitwell Analzer software (Multichannel Systems), and a custom-made MATLAB (The Mathworks, Natrick) code to extract parameters characterizing network activity. Mean firing rate (MFR) was calculated as the average of the spike frequency of all channels across one MEA well. From the burst detection, the number of bursting channels (above threshold 0.4 burst/s and at least 5 spikes in burst with a minimal inter-burst-interval of 100 ms) was determined. A network burst was defined when at least 50% of the channels in one well exhibited a synchronous burst.

### Chemicals

All reagents were prepared fresh into concentrated stocks, and stored frozen at -20 °C. The following compounds were used in pharmacological experiments: Picrotoxin (100 mM in DMSO, Tocris 1128); Forskolin (12 mM in DMSO, Sigma F6886); SQ22536 (50 mM in DMSO, Sigma S153); H89 (5 mM in MQ, Tocris 2910); KT5720 (1 mM in DMSO, Tocris 1283). For all experiment on MEA, before adding chemical to the cultures, an aliquot of the concentrated stock was first diluted in DPBS at room temperature. Then, the appropriate amount of working dilution was added directly to wells on the MEA and mixing was primarily through diffusion into the (500 μl) cell culture medium.

### Pharmacological experiment

Control and *SETD1A^+/-^* networks on MEA were treated with Picrotoxin (PTX, 100 μM), Forskolin (1 μM), H89 (2 μM) and KT5720 (1 μM) at DIV49 or DIV51, and SQ22536 (100 μM) at DIV42 after a 20 min recording of spontaneous activity. Then the recording was stopped temporarily, and the compounds were added to the MEA. We recorded neuronal network activity for 10 min after 5 min treatment of PTX, 60 min treatment of Forskolin, KT5720, H89 and 8 days treatment of SQ22536.

### Whole cell patch clamp

Whole cell patch clamp was performed as previously described (Mossink et al., 2021). Coverslips were placed in the recording chamber of the electrophysiological setup, continuously perfused with oxygenated (95% O_2_/ 5% CO_2_) ACSF at 32°C containing (in mM) 124 NaCl, 1.25 NaH_2_PO4, 3 KCl, 26 NaHCO_3_, 11 Glucose, 2 CaCl_2_, 1 MgCl_2_. Patch pipettes with filament (ID 0.86 mm, OD1.05 mm, resistance 6-8 MΩ) were pulled from borosilicate glass (Science Products GmbH, Hofheim, Germany) using a Narishige PC-10 micropipette puller (Narishige, London, UK). For all recordings of intrinsic properties and spontaneous activity and mPSC activity, a potassium-based intracellular solution containing (in mM) 130 K-Gluconate, 5 KCl, 10 HEPES, 2.5 MgCl_2_, 4 Na_2_-ATP, 0.4 Na_3_-ATP, 10 Na-phosphocreatine and 0.6 EGTA was used, with a pH of 7.2 and osmolality of 290 mOsmol/L. Resting membrane potential (Vrmp) was measured immediately after generation of a whole-cell configuration. Further analysis of active and passive membrane properties was conducted at a holding potential of -60 mV. Passive membrane properties were determined via voltage steps of -10 mV. Active intrinsic properties were measured with a stepwise current injection protocol. Spontaneous postsynaptic currents (sPSCs) and miniature postsynaptic currents (mPSCs) were measured by 10 min continuous recording at a holding potential (V_h_) of -60 mV. In our in-vitro cultures GABAergic spontaneous and miniature events are very sparse (Mossink et al., 2021), thus detected PSCs were considered as mainly reflecting glutamatergic excitatory postsynaptic currents (EPSCs). sEPSC burst inputs were manually counted using clampfit 10.7. sEPSCs were grouped as a burst if at least 3 consecutive events occurred within 50 ms, with at least one of these events showing an amplitude above 100 pA. For the recording of mEPSCs, 1 μM TTX was added to the recording medium. Recordings were conducted at either DIV 21 and DIV 49 (intrinsic properties and sEPSCs) or only DIV 49 (mEPSCs). Cells were visualized with an Olympus BX51WI upright microscope (Olympus Life Science, PA, USA), equipped with a DAGE-MTI IR-1000E (DAGE-MTI, IN, USA) camera) and a CoolLED PE-200 LED system (Scientifica, Sussex, UK) which aided in fluorescent identification of GABAergic neurons. A Digidata 1440A digitizer and a Multiclamp 700B amplifier (Molecular Devices) were used for data acquisition. Sampling rate was set at 20 kHz (intrinsic properties) or 10 kHz (sEPSCs and mEPSCs) and a lowpass 1 kHz filter was used during recording. Recordings were not corrected for liquid junction potential (±10 mV). Recordings were discarded if series resistance reached >25 MΩ or dropped below a 10:0 ratio of membrane resistance to series resistance. Intrinsic electrophysiological properties were analyzed using Clampfit 11.2 (molecular devices, CA, USA), and sEPSCs were analyzed using MiniAnalysis 6.0.2 (Synaptosoft Inc, GA, USA) as previously described (Mossink et al., 2021). Regarding the analysis of the intrinsic properties: In brief, the adaptation ratio was defined as the Δt action potential 8-9/Δt action potential 2-3. The afterhyperpolarization time was defined as the time from which the repolarization phase reaches the threshold potential to the time at which the most hyperpolarized potential was reached. Action potential half-time was calculated as the time to reach 50% ΔmV of the action potential to the AHP peak.

### Calcium imaging

For calcium imaging, cultures were incubated with 4 μg/ml Fluo-8-AM for 30 min at 37°C. After incubation, we removed the excess dye by washing the cells 3 times with HHBS. The cells were then left in culture medium to recover for 15 min. To image the cells, we placed them under the microscope (SliceScope Pro 2000, Scientifica). We continuously perfused the recording chamber with oxygenated (95% O_2_/ 5% CO_2_) and artificial cerebrospinal fluid (ACSF) that was composed of (in mM) 124 NaCl, 3 KCl, 1.25 NaH_2_PO_4_, 2 CaCl_2_, 1 MgCl_2_, 26 NaHCO_3_, 10 Glucose and heated to 37°C. Imaging was performed using a sCMOS camera (Prime BSI Express, Teledyne Photometrics) controlled by Micro-Manager acquisition software (NIH). Fluo-8-AM in the cells was excited at 470 nm by LED (KSL470, Rapp OptoElectronic). We recorded the cells for 2 min with frame rate of 10 Hz. After recording, we analysed the video using MATLAB (The Math Works, Inc. MATLAB. Version 2020b) with a home-made script based on Sun et al (Sun and Südhof, 2020). Circular ROIs were chosen by hand at the centre of the soma with a diameter of 10 μm. We obtained the fluorescent change over time which is defined as ΔF/F = (F-F_0_)/F_0_. Furthermore, the decay of baseline intensity due to bleaching was corrected by exponential fitting to the baseline. Additionally, the traces of each location were analysed for a synchronous firing rate among the selected ROIs to determine network patterns.

### RNA sequencing

Cells were harvested on DIV49 of neuronal differentiation. For RNA-seq, the prepared samples were sequenced on an Illumina NovaSeq SP platform at an average depth of ~50 million reads per sample using a read length of 100 base pairs and an insert size of 350 base pairs. Three biological replicates of control and *SETD1A^+/-^* E/I networks and glutamatergic networks respectively (12 samples in total) using the NucleoSpin RNA isolation kit (Machery Nagel, 740955.250) according to the manufacturer’s instructions. RNA yield was quantified with a NanoDrop Spectrophotometer (NanoDrop Technologies, Wilmington, DE, USA) and RNA integrity was assessed with Bioanalyzer 2100 RNA 6000 Nano Kit (Agilent Technologies, Santa Clara, CA, USA). All samples had an RNA Integrity Number (RIN) > 9. Library preparation and paired-end RNA-sequencing were carried out at the Norwegian High-Throughput Sequencing Centre (www.sequencing.uio.no). Briefly, libraries were prepared with the TruSeq Stranded mRNA kit from Illumina which involves Poly-A purification to capture coding as well as several non-coding RNAs. The prepared samples were then sequenced on an Illumina NovaSeq SP platform.

### RNA-seq data processing

Raw sequencing reads were quality assessed with FastQC (Babraham Institute). To pass the initial QC check, the average Phred score of each base position across all reads had to be at least 30. Reads were further processed by cutting individual low-quality bases and removing adapter and other Illumina-specific sequences with Trimmomatic V0.32 using default parameters (Bolger et al., 2014). The trimming process may result in some reads being discarded and their mates thereby unpaired, therefore only reads that remained paired after trimming were used for downstream analyses. Since the cultures contained both hiPSC-derived neurons as well as rat astrocytes which were added to support neuronal maturation, sequencing reads were separated according to their species of origin using the *in silico* RNA-seq read sorting tool Sargasso (Qiu et al., 2018), which was able to successfully eliminate sequencing reads stemming from rat astrocytes (**Figure S12**). To quantify gene expression levels, reads mapped by Sargasso were summarized at the gene level using featureCounts (Liao et al., 2014) guided by ENSEMBL annotations.

### Differential expression (DE) analysis and over-representation test

To evaluate the species separation performance of Sargasso, bioinformatical estimation of cell type abundances (deconvolution) was carried out with CIBERSORTx (Newman et al., 2019) using expression signatures for human neurons and rodent astrocytes (Zhang et al., 2016) (**Figure S12**). Before conducting the DE analyses, genes with very low to zero expression were removed by filtering out any gene with ≤1 counts per million (CPM) in 3 or more samples (the smallest group size). DE analysis was performed using the statistical R package DESeq2 (Love et al., 2014), which provides methods to test for differentially expressed genes by use of negative binomial generalized models. The DESeq2 workflow begins by taking raw read count data as input and applies an internal normalization method that corrects for sequencing depth and RNA composition. The standard DE analysis consists of size factor estimation, dispersion estimation, and model fitting, as well as an independent filtering step that optimizes the number of significant DE genes. After the pre-filtering and independent filtering steps, a total of 14,144 genes were retained and examined in the DE analyses. A DE gene was considered significant if the FDR was <0.05. Gene Ontology (GO) enrichment tests of significant DE gene sets were conducted with the over-representation analysis tool clusterProfile (Yu et al., 2012) using the enrichGO function. A GO term was considered significantly enriched if the FDR was <0.05. Disease association analysis was performed with the R package disgenet2r (Piñero et al., 2020) using both the CURATED database, which includes more than 10,000 somatic and mental disorders, and the PSYGENET, which includes 109 mental diseases. In both cases, FDR<0.05 was used as the threshold to determine significant disease association.

### Data analysis

The statistical analysis of the data were performed using GraphPad Prism 8 (GraphPad Software, Inc., CA, USA). We ensured normal distribution using a Shapiro-Wilk normality test. Analysis was done using unpaired Student’s t tests when comparing two variables at a single time point, or one-way ANOVA with sequential post hoc Bonferroni corrections. Results with p values lower than 0.05 were considered as significantly different. p <0.05 (*), p <0.01 (**), p <0.001 (***). Data is shown as mean ± standard error of the mean (SEM).

## Data availability

All the data are available within this manuscript and its Supplementary information. RNA-seq raw and processed data are available from GEO repository (GEO: GSE180648). Additional information is available from the corresponding author upon reasonable request.

## Acknowledgements

This work was supported by grants from: ERA-NET NEURON-102 SYNSCHIZ grant (NWO) 013-17-003 4538 (to D.S) and ERA-NET NEURON DECODE! grant (NWO) 013.18.001 (to N.N.K); The Netherlands Organization for Scientific Research, NWO-CAS grant 012.200.001 (to N.N.K); the Netherlands Organization for Health Research and Development ZonMw grant 91217055 (to. H.v.B and N.N.K); SFARI grant 610264 (to N.N.K); Marie Curie Actions European Fellowship 794273 (to N.K); South-Eastern Norway Regional Health Authority #2018094 (to S.D); China scholarship council 201806210076 (to S.W).

## Author contribution

S.W, D.S and N.N.K conceived and designed all the experiments. D.S, N.N.K and H.v.B supervised the study. S.W, J.v.R, I.A, N.K, N.M, A.B, E.L, K.W, C.S performed all experiments. S.W, J.v.R, I.A, N.K, N.M, A.B, D.S, N.N.K performed data analysis. S.W, D.S, J.v.R, I.A and N.N.K wrote the manuscript. H.v.B, S.D and T.K edited the manuscript.

## Declaration of interests

The authors declare no competing interests.

## Notes

### Competing Interest Statement

The authors have declared no competing interest.

