## Supplementary files for "Loss-of-function variants in the schizophrenia risk gene *SETD1A* alter neuronal network activity in human neurons through cAMP/PKA pathway"

Dept. Cognitive Neuroscience

Donders Institute for Brain, Cognition and Behaviour

Radboud University Nijmegen Medical Centre

Postbus 9101

6500 HB Nijmegen

The Netherlands

**Supplementary Figures**

- **Supplementary figure S1**: Characterization of control and *SETD1A*$^{+/-}$ iPSCs.
- **Supplementary figure S2**: Related to Figure 1.
- **Supplementary figure S3**: Altered network activity in *SETD1A*$^{+/-}$ neurons recorded by calcium imaging.
- **Supplementary figure S4**: Related to Figure 1.
- **Supplementary figure S5**: Related to Figure 2.
- **Supplementary figure S6**: *SETD1A* haploinsufficiency in glutamatergic neurons leads to alteration of transcriptomic profile.
- **Supplementary figure S7**: Related to Figure 4.
- **Supplementary figure S8**: Related to Figure 4.
- **Supplementary figure S9**: Related to Figure 5.
- **Supplementary figure S10**: Related to Figure 5.
- **Supplementary figure S11**: Related to Figure 5.
- **Supplementary figure S12**: Separation of mixed-species RNA-seq reads according to species of origin.

**Supplementary Videos**

- Related to supplementary figure S3.

**Supplementary Tables**

- **Supplementary table S1**: Cell type specific electrophysiological properties of neurons in E/I cultures during development.
- **Supplementary table S2**: Overrepresented DEGs in “Schizophrenia” from disease enrichment analysis using DisGeNET database.
- **Supplementary table S3**: Upregulated DEGs involve in synaptic strength.
- **Supplementary table S4**: Comparison of DEGs with transcriptomic profile of the published *Setd1a*$^{+/-}$ mice.
- **Supplementary table S5**: Comparison of DEGs with SFARI datasets.
- **Supplementary table S6**: Cell type specific electrophysiological properties of neurons in glutamatergic cultures during development.
- **Supplementary table S7**: Upregulated DEGs involve in second messenger signaling. Related to Figure 5a.

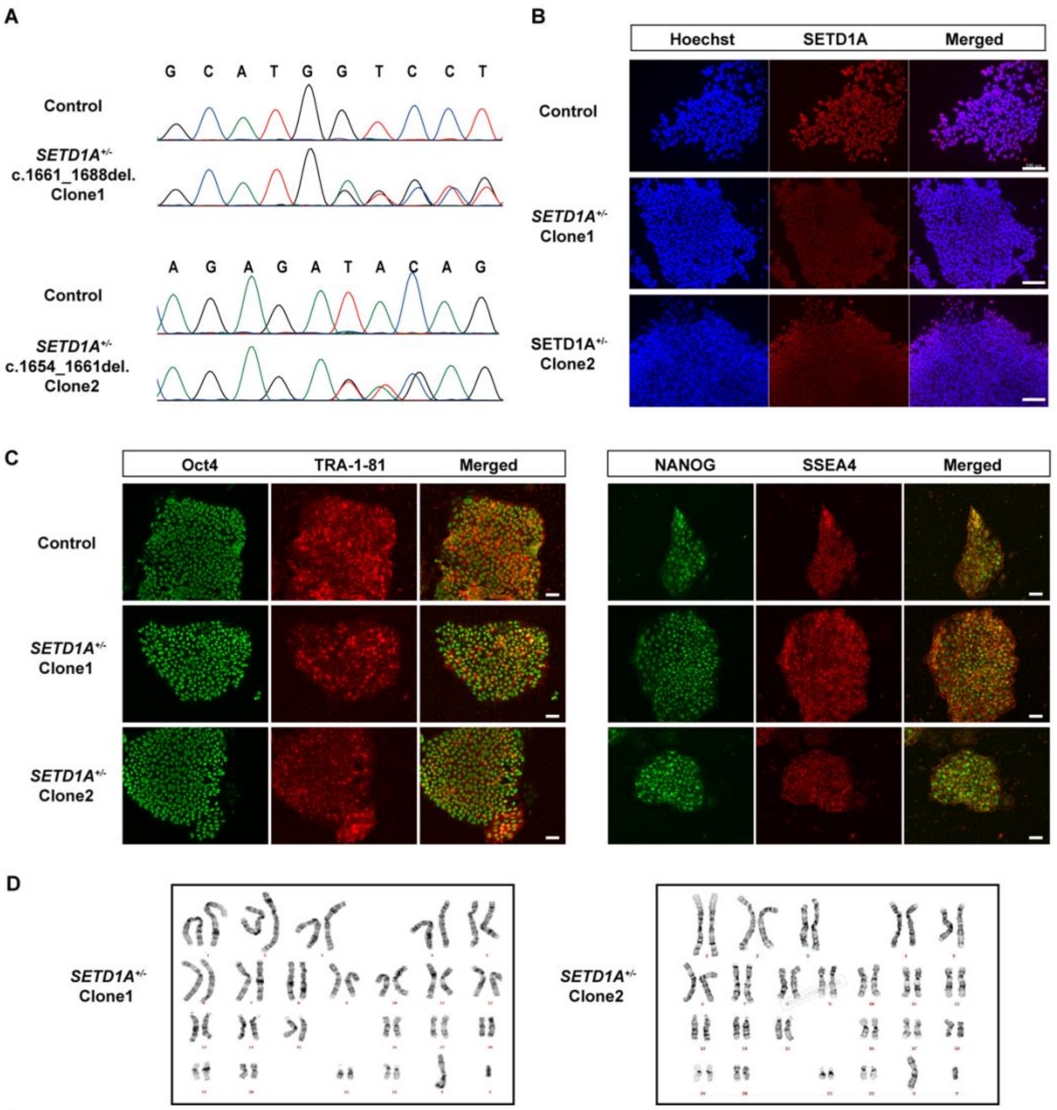


E

| Gene | Chr | Position | Strand | Sequence | # mismatch | Score | On target | Primers |
|---|---|---|---|---|---|---|---|---|
| LINC00485 | 16 | 103212012 | -1 | CCTGTATCTCTGGCCTCAGGTAC | 3 | 0.86 | False | Fw: ATCCCCACATGTGCAGTGAG<br>Rv: TAGTCCTCAAGAGTCTGGGCT |
| None | 1 | 48188845 | -1 | CCTGTTTCTCTGGCCCCAAGGTC | 2 | 0.56 | False | Fw: CGAGGAGGTTGGGGTGTCAAG<br>Rv: CGCTCCAACATATGCTGTGC |
| KIAA1671 | 22 | 25388422 | -1 | CCTGTATCTCGGCTCCTAGACAG | 3 | 0.48 | False | Fw: CAGGGTCTCAGATGGAGGGA<br>Rv: TCCAGGTGATAACTGCTGGC |
| BNIP | 5 | 172590193 | -1 | CCAGTATCCCTGGCCTGAGGGA | 4 | 0.44 | False | Fw: GTTCTGACCTCCAGCTGACC<br>Rv: CAGCCAAATGCACCAGAACC |
| PCLO | 7 | 82581887 | 1 | GTCTCAGTGGCCAGAGATACAGG | 4 | 043 | False | Fw: TTCCCTGGGATACGGTGCTA<br>Rv: TCAAGCAGCTCCCACATCAG |

**Supplementary figure S1. Characterization of control and *SETD1A*$^{+/-}$ iPSCs. (a)** Sanger sequence of control and *SETD1A*$^{+/-}$ iPSCs. **(b)** Representative images showing reduced expression of SETD1A in *SETD1A*$^{+/-}$ iPSCs. Scale bar = 100 μm. **(c)** Representative images of control and *SETD1A*$^{+/-}$ iPSCs stained for pluripotent markers. Scale bar = 50 μm. **(d)** *SETD1A*$^{+/-}$ iPSCs maintained a normal karyotype. **(e)** Top five potential off-target sites have been sequenced and no mutations were detected.

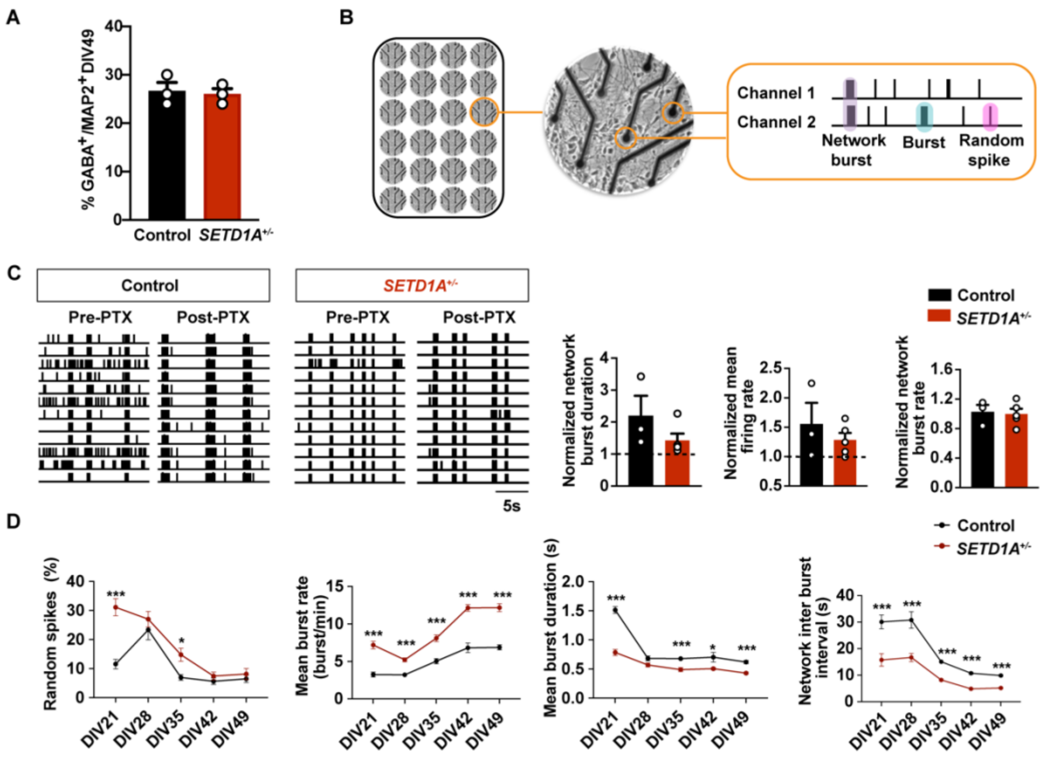


**Supplementary figure S2. *SETD1A*$^{+/-}$ neurons exhibit dysregulated neuronal network organization, Related to Figure 1. (a)** Quantification of MAP2 and GABA co-localized neurons in networks derived from control and *SETD1A*$^{+/-}$ iPSCs at DIV49. **(b)** Schematic overview of an electrophysiological recording from neurons cultured on Micro-electrode arrays (MEAs). **(c)** Representative raster plots of 20 sec showing the effect of acute treatment of 100 μM picrotoxin on control and *SETD1A*$^{+/-}$ E/I networks at DIV49; Quantification of network burst duration, mean firing rate and network burst rate normalized to their respective baseline recording. n = 3 individual wells for control. n = 5 individual wells for *SETD1A*$^{+/-}$. **(d)** Quantification of network parameters using MEA recording as indicated. n = 40 individual wells for control. n = 63 individual wells for *SETD1A*$^{+/-}$.

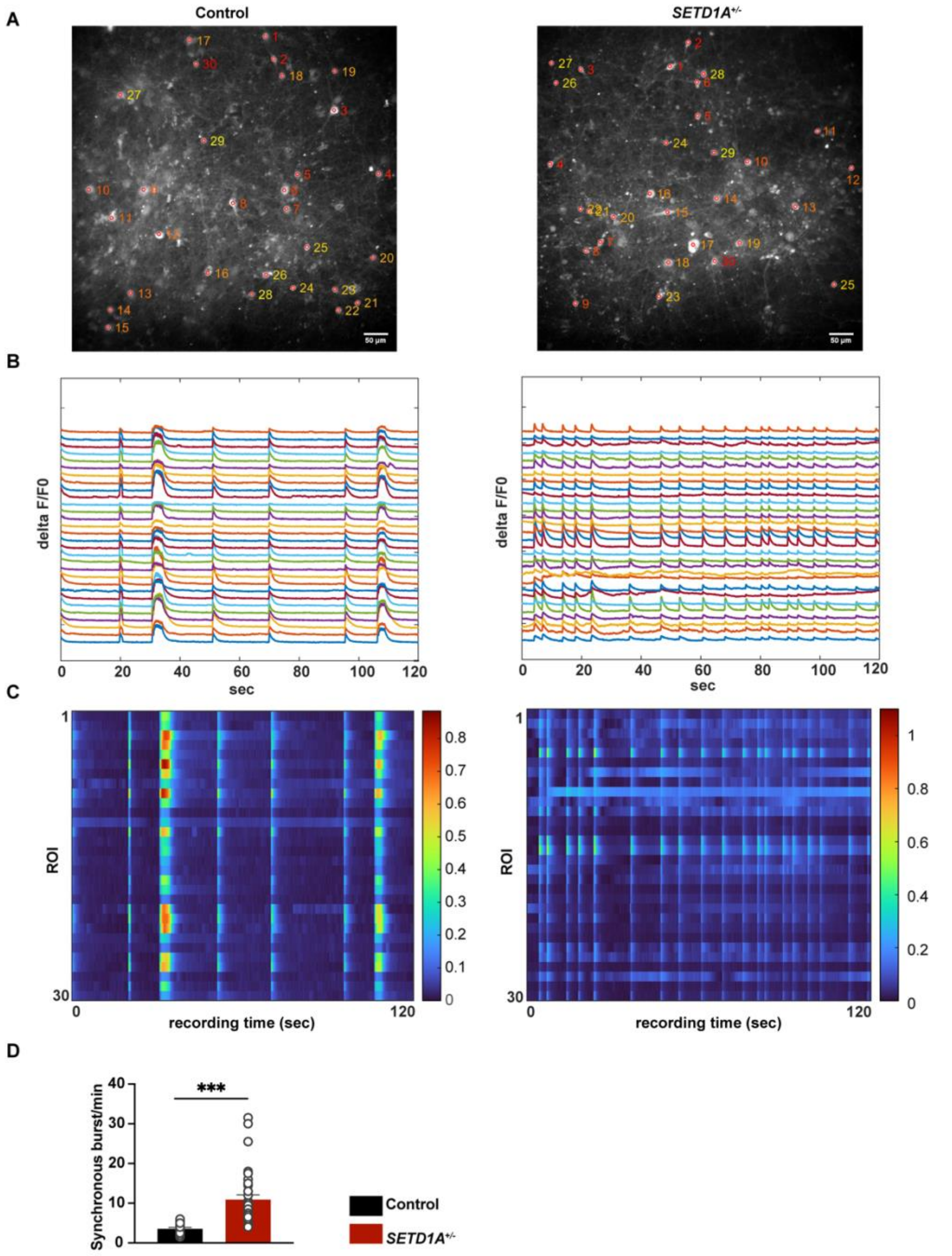


**Supplementary figure S3. Altered network activity in *SETD1A*$^{+/-}$ neurons recorded by calcium imaging. (a)** Representative ROIs in the field of view showing calcium fluctuations in control (left) and *SETD1A*$^{+/-}$ (right) neurons at DIV70. Scale bar = 50 μm. Representative videos can be seen from Supplementary Video1 for control and Supplementary Video 2 for *SETD1A*$^{+/-}$. **(b-c)** Changes in calcium signal from 30 neurons showed by temporal calcium traces (b) and heatmap (c) in a 2 min recording from control (left) and *SETD1A*$^{+/-}$ (right) neurons. **(d)** Quantification of synchronous burst rate. n = 23 for control. n = 34 for *SETD1A*$^{+/-}$. *p < 0.05, **p < 0.01, ***p < 0.001, unpaired Student's t test was performed between control and *SETD1A*$^{+/-}$ cultures.

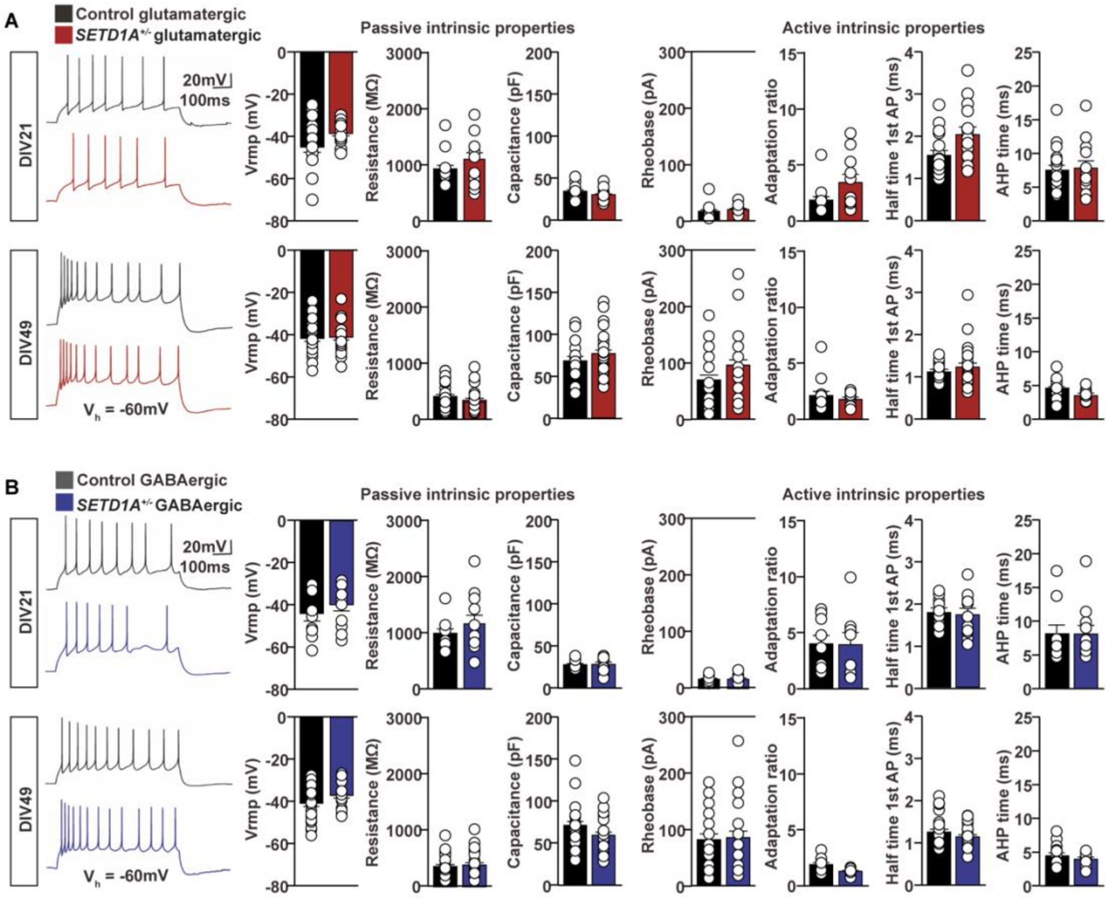


**Supplementary figure S4. Basic passive and active intrinsic electrophysiological properties of control and *SETD1A*$^{+/-}$ neurons in E/I networks during development, Related to Figure 1. (a)** Glutamatergic neurons: Representative whole-cell patch-clamp recordings of action potential firing pattern and quantitative analyses of basic intrinsic properties of control and *SETD1A*$^{+/-}$ at DIV 21 and DIV49. DIV 21: Control (passive properties n = 18, active properties n = 15), *SETD1A*$^{+/-}$ (passive properties n = 15, active properties n = 11). DIV 49: Control (passive properties n = 28, active properties n = 15), *SETD1A*$^{+/-}$ (passive properties n = 27, active properties n = 12).
**(b)** GABAergic neurons: Representative whole-cell patch-clamp recordings of action potential firing pattern and quantitative analyses of basic intrinsic properties of control and *SETD1A*$^{+/-}$ glutamatergic neurons at DIV 21 and DIV49. DIV 21: Control (passive properties n = 11, active properties n = 9), *SETD1A*$^{+/-}$ (passive properties n = 11, active properties n = 8). DIV 49: Control (passive properties n = 27, active properties n = 14), *SETD1A*$^{+/-}$ (passive properties n = 27, active properties n = 11). Groups were

compared using Students' T-test with Bonferroni correction for multiple testing. Except for the resting membrane potential (Vrmp), parameters were determined at a holding potential ($V_h$) of -60 mV. Action potential (AP), after-hyperpolarization (AHP).

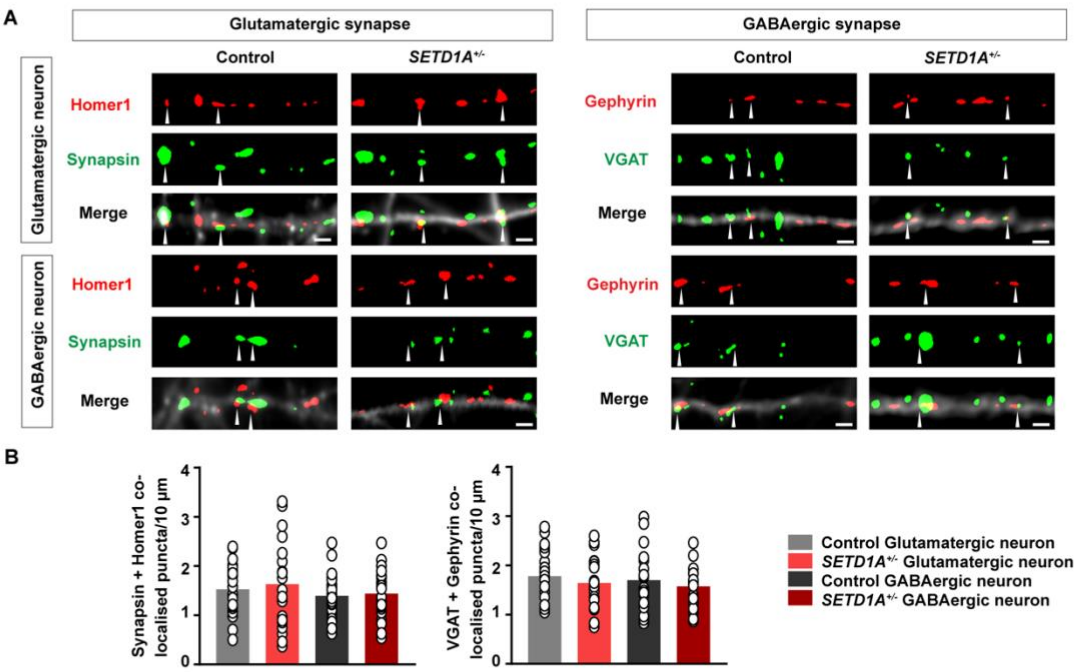


**Supplementary figure S5**: ***SETD1A* haploinsufficiency shows no effect on glutamatergic and GABAergic synapse density on each cell type, Related to Figure 2. (a)** Representative images of immunocytochemistry stained for glutamatergic synapse (Synapsin as a presynaptic marker and Homer1 as a postsynaptic marker) and GABAergic synapse (VGAT as a pre-synaptic marker and Gephyrin as a post synaptic marker) at DIV49 on each cell type. White arrows indicate the co-localized Synapsin/Homer1 or VGAT/Gephyrin puncta. Scale bar = 2 μm **(b)** Quantification of the density of co-localized Synapsin/Homer1 and VGAT/Gephyrin puncta on glutamatergic neurons (Synapsin/Homer1: n = 22 for control, n = 23 for *SETD1A*$^{+/-}$, VGAT/Gephyrin: n = 22 for control, n = 22 for *SETD1A*$^{+/-}$) and GABAergic neurons (Synapsin/Homer1: n = 20 for control, n = 20 for *SETD1A*$^{+/-}$, VGAT/Gephyrin: n = 20 for control, n = 20 for *SETD1A*$^{+/-}$).

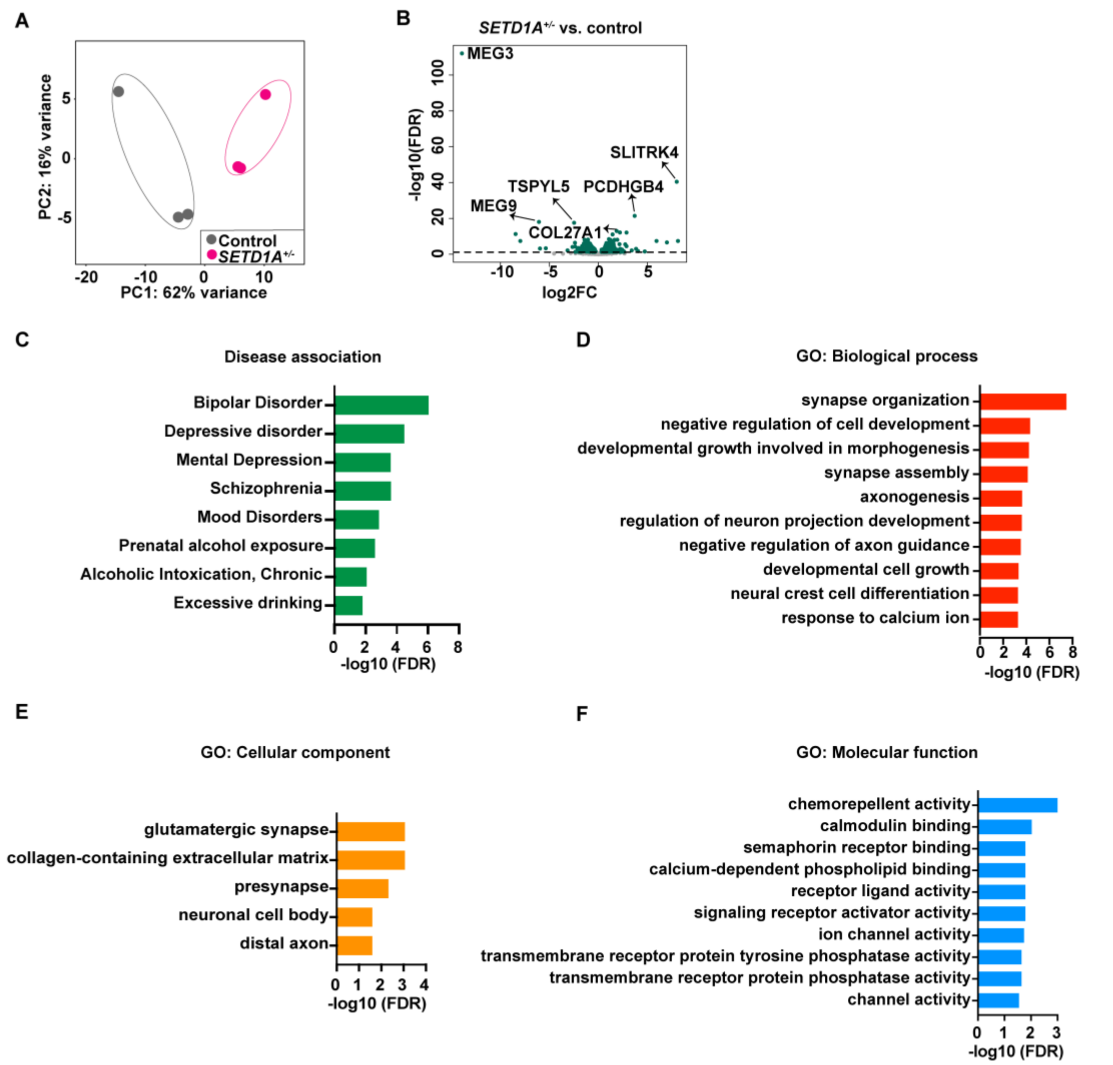


**Supplementary figure S6. *SETD1A* haploinsufficiency in glutamatergic neurons leads to alteration of transcriptomic profile. (a)** PCA showing tight clustering of 3 replicates for each genotype. **(b)** Volcano plots of -log10 (FDR) versus the log2 (fold change) of transcript levels for all genes. Relative to control, significantly up or down-regulated genes are shown in green. Top 3 upregulated and downregulated genes are labeled. **(c)** Disease terms of DisGeNET database associated with differentially expressed genes (DEGs). **(d-f)** Gene Ontology (GO) term analysis of DEGs.

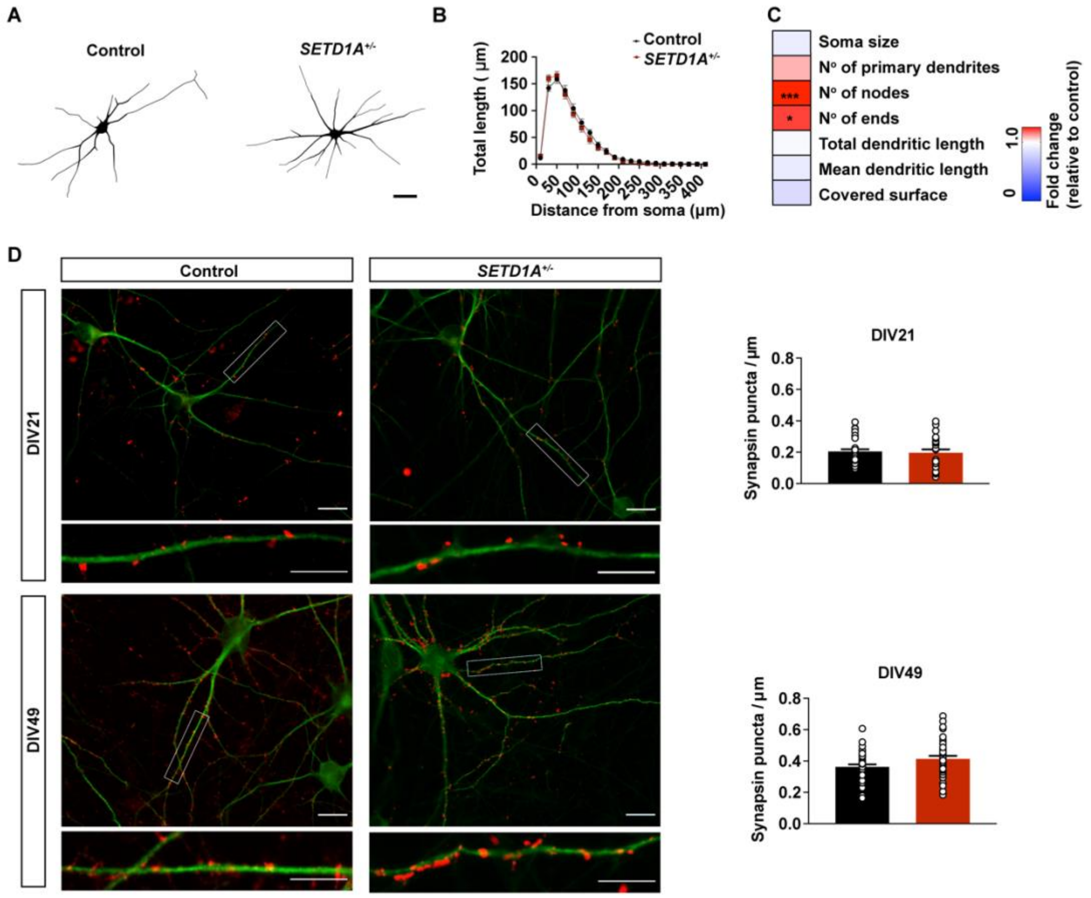


**Supplementary figure S7. *SETD1A* haploinsufficiency in glutamatergic neurons shows no effect on Synapsin density, Related to Figure 4. (a-b)** Representative somatodendritic reconstructions of glutamatergic neurons and Sholl analysis in control and *SETD1A*$^{+/-}$ networks at DIV21. (sample size: n = 45 for control glutamatergic neurons. n = 49 for *SETD1A*$^{+/-}$ glutamatergic neurons) Scale bar = 40 µm. Data represent means ± SEM. *p < 0.05, **p < 0.01, ***p < 0.001, two-way ANOVA with post hoc Bonferroni correction. **(c)** Heatmap showing fold change of all the parameters compared to control in reconstruction for glutamatergic neurons at DIV21. *p < 0.05, **p < 0.01, ***p < 0.001, unpaired Student's t test. **(d)** Representative images of control and *SETD1A*$^{+/-}$ glutamatergic neurons stained for MAP2 (green) and Synapsin (red) at DIV21 and DIV49 (Scale bar = 20 µm) and quantification of Synapsin puncta per µm. (sample size: n = 41 for control glutamatergic neurons. n = 35 for *SETD1A*$^{+/-}$ glutamatergic neurons at DIV21; n = 37 for control glutamatergic neurons. n = 36 for *SETD1A*$^{+/-}$ glutamatergic neurons at DIV49).

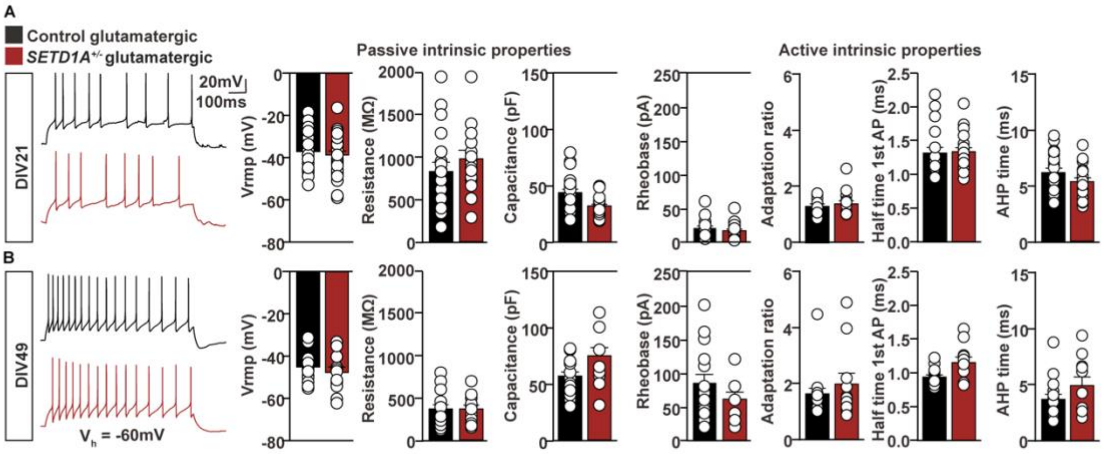


**Supplementary figure S8.** ***SETD1A*** **haploinsufficiency in glutamatergic neurons shows no effect on intrinsic properties, Related to Figure 4. (a)** Intrinsic properties of Glutamatergic neurons in glutamatergic cultures at DIV 21. Control/*SETD1A*$^{+/-}$ n = 33/32 (passive properties) and 18/20 (active properties). **(b)** Intrinsic properties of glutamatergic neurons in glutamatergic cultures at DIV 49. Control/*SETD1A*$^{+/-}$ n = 17/12 (passive properties) and 16/11 (active properties). Groups were compared using Students' T-test with Bonferroni correction for multiple testing. Except for the resting membrane potential ($V_{rmp}$), parameters were determined at a holding potential ($V_h$) of -60 mV. Action potential (AP), after-hyperpolarization (AHP).

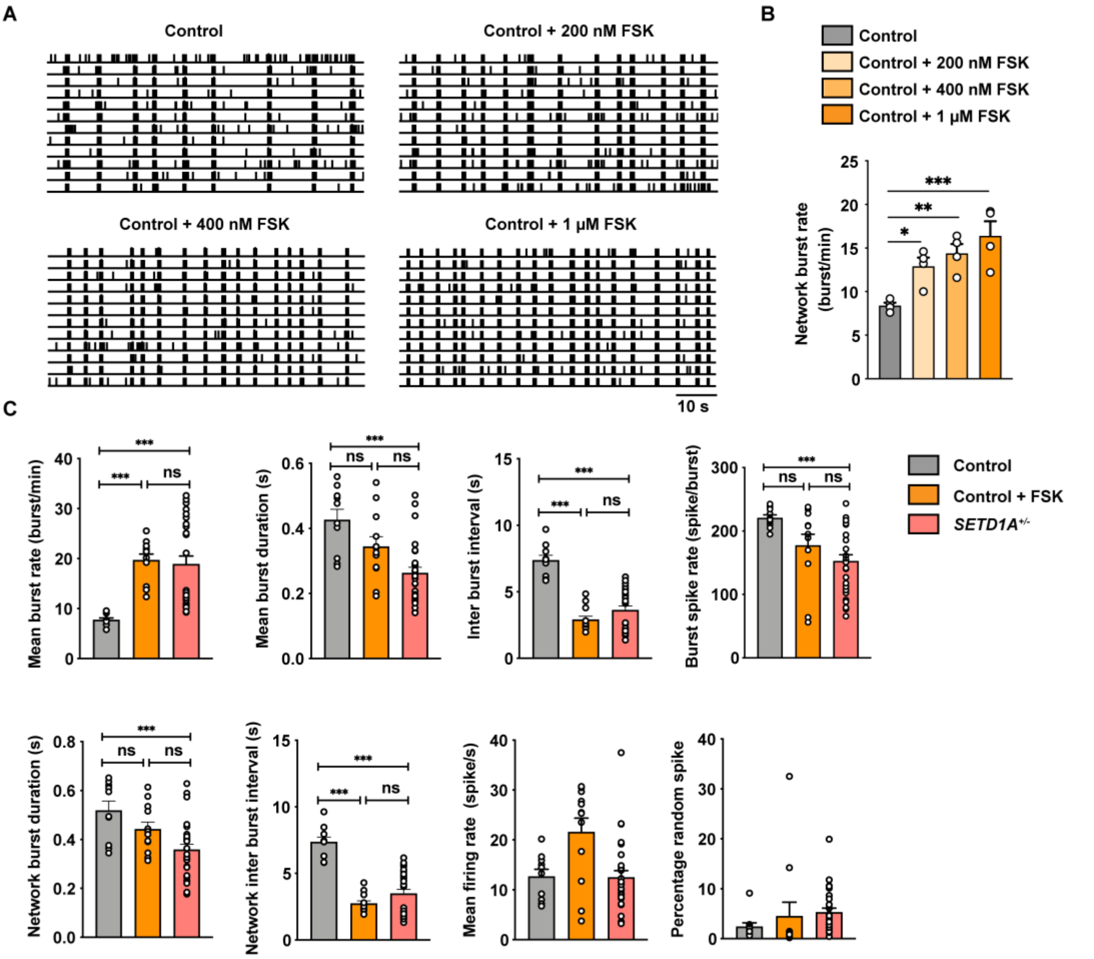


**Supplementary figure S9. Control networks treated with Forskolin mimic the phenotype of *SETD1A*$^{+/-}$ networks, Related to Figure 5. (a-b)** Representative raster plot for 1 min and quantification of network burst rate showing the dose-dependent effect of AC agonist forskolin on control networks (sample size: control n= 4 wells, control + forskolin n = 4 wells) **(c)** Control networks were treated with 1 μM forskolin for 1 hour. Quantification of network parameters recorded by MEA as indicated. (sample size: control n =10 wells, control + forskolin n = 12 wells, *SETD1A*$^{+/-}$ n = 28 wells). *P <0.05, **P < 0.01, ***P < 0.001, one-way ANOVA test with post hoc Bonferroni correction.

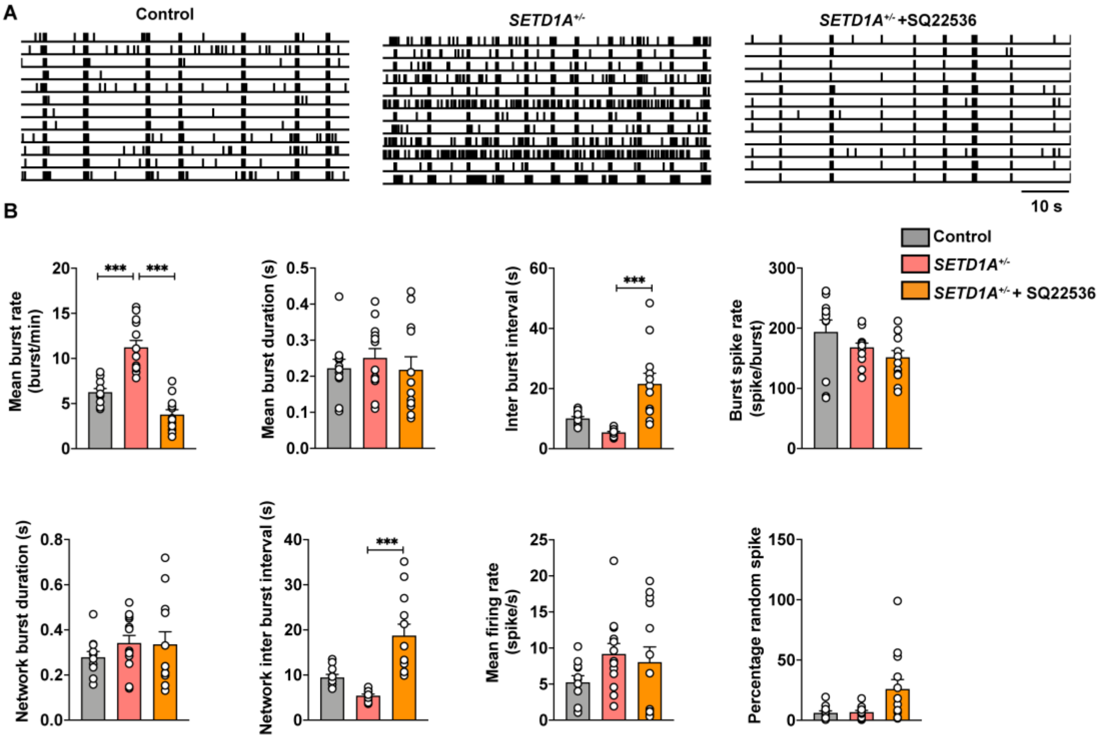


**Supplementary figure S10. The phenotype of *SETD1A*$^{+/-}$ networks can be rescued by SQ22536, Related to Figure 5. (a-b)** Representative raster plot for 1 min and quantification of network burst rate showing the effect of adenylate cyclase inhibitor SQ22536 (100 μM) on *SETD1A*$^{+/-}$ networks (sample size: control n = 11 wells, *SETD1A*$^{+/-}$ + SQ22536 n = 12 wells, *SETD1A*$^{+/-}$ n = 13 wells). *P <0.05, **P < 0.01, ***P < 0.001, one-way ANOVA test with post hoc Bonferroni correction.

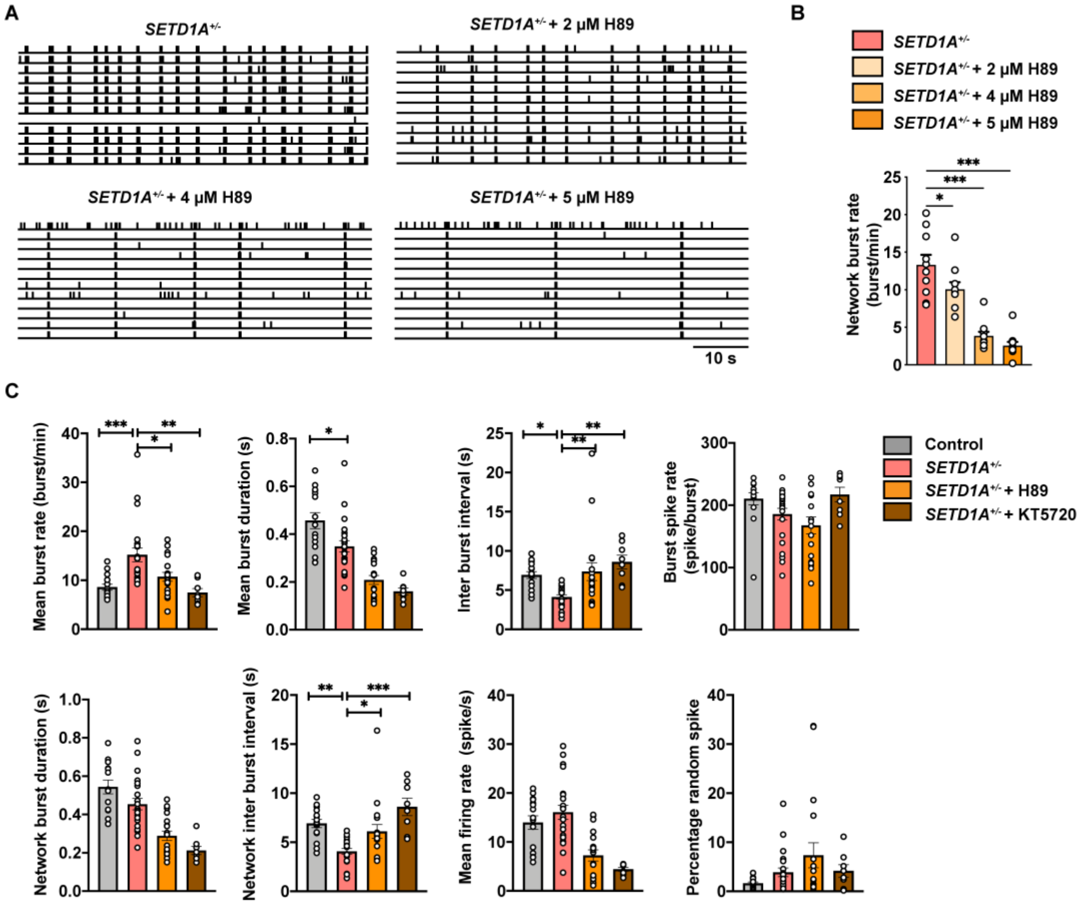


**Supplementary figure S11. The phenotype of *SETD1A*$^{+/-}$ networks can be rescued by H89 or KT5720, Related to Figure 5. (a, b)** Representative raster plot for 1 min and quantification of network burst rate showing the dose-dependent effect of PKA inhibitor H89 on *SETD1A*$^{+/-}$ network. (sample size: *SETD1A*$^{+/-}$ n = 10 wells, *SETD1A*$^{+/-}$ + H89 n = 10 wells) **(c)** *SETD1A*$^{+/-}$ networks were treated with 2 μM H89 or 1 μM KT5720 for 1 hour. Quantification of network parameters recorded by MEAs as indicated. (sample size: control n = 15 wells, *SETD1A*$^{+/-}$ + H89 n = 18 wells, *SETD1A*$^{+/-}$ + KT5720 n = 18 wells, *SETD1A*$^{+/-}$ n = 22 wells). Data represent means ± SEM. *P < 0.05, **P < 0.01, ***P < 0.001, one-way ANOVA test with post hoc Bonferroni correction.

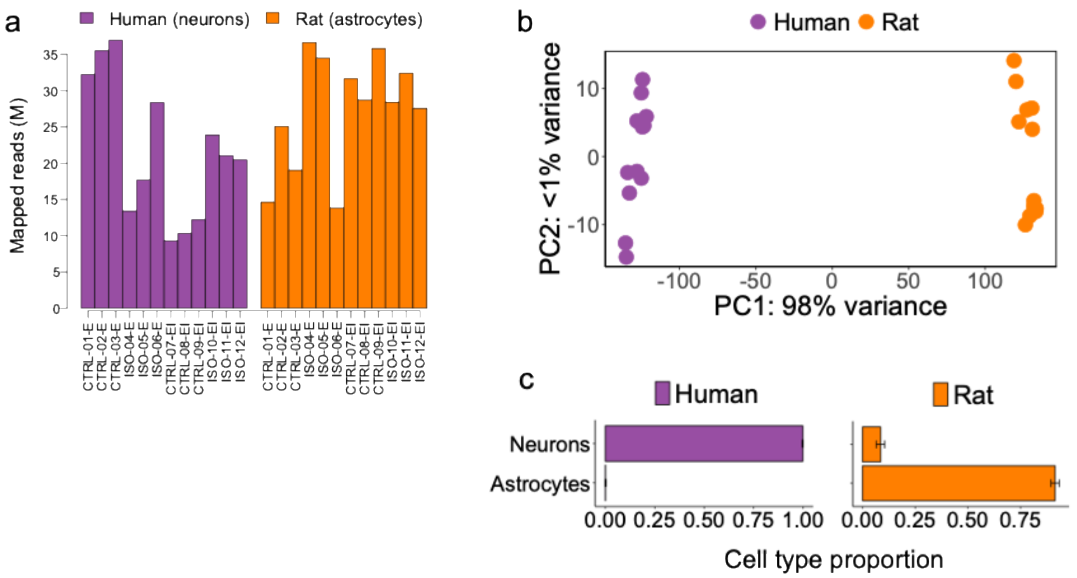


**Supplementary figure S12. Separation of mixed-species RNA-seq reads according to species of origin. (a)** Mapped reads (in millions) per sample per species after bioinformatically separating human neuronal reads from rat astrocytic reads. Average sequencing depth per sample (sum of rat and human reads) was 49.1 million reads. **(b)** PCA plot of samples showing complete separation of clusters according to species of origin. Top 1000 genes with largest variance across samples were used as input. **(c)** Deconvolution of cell type proportions showing no contamination of astrocytic reads in human samples after species separation.

**Supplementary table S1:** Cell type specific electrophysiological properties of neurons in E/I cultures during development. Statistical testing compared genotypes per cell class and DIV.* = based on KS-test on cumulative data.

| **E/I co-culture** | **Glutamatergic neurons** | | | **GABAergic neurons** | | |
|---|---|---|---|---|---|---|
| **DIV 21** | Control | *SETD1A*$^{+/-}$ | p-value | Control | *SETD1A*$^{+/-}$ | p-value |
| **Passive intrinsic properties** | n = 18 | n = 15 | | n = 27 | n = 27 | |
| Resting membrane potential (mV) | -45.1 ± 3.1 | -38.5 ± 1.3 | | -46.6 ± 3.6 | -42.1 ± 3.1 | |
| Membrane resistance (MΩ) | 976 ± 68 | 1157 ± 123 | | 1044 ± 84 | 1223.0 ± 161 | |
| Membrane capacitance (pF) | 36.7 ± 1.8 | 32.2 ± 2.0 | | 28.9 ± 1.5 | 29.3 ± 2.7 | |
| **Active intrinsic properties** | (n = 15) | (n = 11) | p-value | (n = 14) | (n = 11) | p-value |
| AP threshold (mV) | -27.9 ± 1.1 | -25.3 ± 1.4 | | -26.6 ± 1.3 | -27.0 ± 1.4 | |
| Rheobase (pA) | 18.1 ± 2.7 | 21.3 ± 2.3 | | 17.5 ± 1.9 | 17.7 ± 2.3 | |
| 1st AP amplitude | 98.4 ± 2.2 | 98.7 ± 3.4 | | 99.2 ± 2.6 | 99.6 ± 4.8 | |
| 1st AP Half time (ms) | 1.6 ± 0.1 | 2.0 ± 0.2 | | 1.9 ± 0.1 | 1.8 ± 0.2 | |
| Adaptation rate 8-9rd/2-3rd AP | 2.0 ± 0.3 | 3.6 ± 0.7 | | 4.2 ± 0.8 | 4.1 ± 1.1 | |
| 1st AP AHP time (ms) | 8.0 ± 0.8 | 8.3 ± 1.2 | | 8.6 ± 1.4 | 8.6 ± 1.3 | |
| 1st-2nd AP AHP amplitude (mV) | 2.8 ± 0.5 | 0.5 ± 1.4 | | 0.8 ± 1.1 | 0.6 ± 0.9 | |
| **Synaptic inputs/ postsynaptic events** | (n = 23) | (n = 22) | p-value | (n = 21) | (n = 22) | p-value |
| sEPSC amplitude (pA) | 31.0 ± 2.0 | 30.3 ± 2.6 | | 28.9 ± 3.9 | 25.6 ± 1.6 | |
| sEPSC frequency (Hz) | 1.8 ± 0.5 | 1.9 ± 0.4 | | 1.6 ± 0.3 | 1.8 ± 0.6 | |
| sEPSC synchronous inputs (n/min) | 0.5 ± 0.3 | 0.7 ± 0.2 | | 1.9 ± 0.4 | 1.6 ± 0.4 | |

**DIV 49**

| | | | | | | |
|---|---|---|---|---|---|---|
| **Passive intrinsic properties** | (n = 28) | (n = 27) | p-value | (n = 27) | (n = 27) | p-value |
| Resting membrane potential (mV) | -45.3 ± 1.6 | -44.6 ± 1.6 | | -44.4 ± 1.6 | -40.3 ± 1.4 | |
| Membrane resistance (MΩ) | 435.0 ± 44.4 | 361.0 ± 42.9 | | 386.3 ± 38.5 | 415.3 ± 46.2 | |
| Membrane capacitance (pF) | 73.5 ± 4.6 | 82.9 ± 6.0 | | 76.6 ± 5.2 | 64.0 ± 4.1 | |
| **Active intrinsic properties** | (n = 15) | (n = 12) | p-value | (n = 14) | (n = 11) | p-value |
| AP threshold (mV) | -32.0 ± 1.9 | -27.7 ± 1.3 | | -28.8 ± 1.7 | -27.0 ± 1.0 | |
| Rheobase (pA) | 75.0 ± 9.8 | 103.8 ± 13.4 | | 89.0 ± 11.0 | 92.9 ± 12.9 | |
| 1st AP amplitude | 104.5 ± 2.5 | 99.4 ± 3.1 | | 104.6 ± 2.7 | 106.7 ± 2.5 | |
| 1st AP Half time (ms) | 1.2 ± 0.1 | 1.3 ± 0.1 | | 1.3 ± 0.1 | 1.2 ± 0.1 | |
| Adaptation rate 8-9rd/2-3rd AP | 2.3 ± 0.4 | 1.9 ± 0.2 | | 2.0 ± 0.2 | 1.4 ± 0.1 | 0.005 |
| 1st AP AHP time (ms) | 4.9 ± 0.4 | 3.7 ± 0.3 | | 4.7 ± 0.5 | 4.1 ± 0.3 | |
| 1st-2nd AP AHP amplitude (mV) | 1.8 ± 0.3 | 2.2 ± 0.5 | | 1.7 ± 0.7 | 1.7 ± 0.7 | |
| **Synaptic inputs/ postsynaptic events** | (n = 15) | (n = 13) | p-value | (n = 18) | (n = 16) | p-value |
| sEPSC amplitude (pA) | 24.8 ± 2.4 | 22.4 ± 1.0 | | 22.3 ± 1.9 | 24.0 ± 2.5 | |
| sEPSC frequency (Hz) | 2.5 ± 0.7 | 2.2 ± 0.6 | | 1.6 ± 0.3 | 1.6 ± 0.4 | |
| sEPSC synchronous inputs (n/min) | 1.1 ± 0.2 | 2.0 ± 0.4 | 0.031 | 0.8 ± 0.1 | 1.6 ± 0.3 | 0.009 |
| | (n = 17) | (n = 16) | p-value | (n = 16) | (n = 13) | p-value |
| mEPSC amplitude (pA) | 17.3 ± 1.0 | 20.2 ± 1.5 | <0.001* | 16.6 ± 1.2 | 19.5 ± 2.1 | <0.001* |
| mEPSC frequency (Hz) | 1.4 ± 0.3 | 2.0 ± 0E.5 | <0.001* | 1.1 ± 0.3 | 2.8 ± 0.6 | <0.001* |

**Supplementary table S2**: Overrepresented DEGs in “Schizophrenia” from disease enrichment analysis using DisGeNET database.

| List of genes | |
|---|---|
| Schizophrenia | *ABCB1,ADNP,ALDH1A2,APOL2,ASTN2,BACE1,BCL9,CACNG8,CCDC86,CCND2,CDKN1C,CHGA,CHGB,CNIH3,CPLX1,CRH,DCC,DKK3,DLG2,DNMT3B,DRD2,DTNBP1,FABP5,FADS2,FASTKD5,GDNF,GFRA1,GRIK3,GRIN2A,GRM3,GRM4,GSK3A,HDAC2,HLAC,HTR3B,HTR7,KCNB1,KLF12,KMT2A,LPAR1,MAGI1,MAOB,MBP,MET,MYO16,NR3C1,NRXN3,PANK2,PBRM1,PCDH17,PHOX2B,PLCB1,PLXNA2,PNPO,PTPN21,PTPRA,S100B,SLC12A5,SLC25A27,SLC6A2,SLIT3,SNCB,ST3GAL1,SULT4A1,TNFRSF1A,TPI1,UCP2,ZSCAN31,ABCB1,ADNP,ALDH1A2,ASTN2,BACE1,BCL9,CACNG8,CCDC86,CCND2,CDKN1C,CHGA,CHGB,CNIH3,CRH,DCC,DKK3,DLG2,DNMT3B,FABP5,FADS2,GDNF,GFRA1,GRIK3,GRIN2A,GRM4,HDAC2,HLAC,HTR3B,KMT2A,LPAR1,MAGI1,MBP,MYO16,NRXN3,PANK2,PBRM1,PCDH17,PHOX2B,PNPO,PTPN21,PTPRA,S100B,SLC12A5,SLC25A27,SLC6A2,SLIT3,SNCB,ST3GAL1,SULT4A1,TNFRSF1A,TPI1,UCP2,ZSCAN31* |

**Supplementary table S3**: Upregulated DEGs involve in synaptic strength.

| Term | List of genes |
|---|---|
| Neurotransmitter transport | *DTNBP1,MAOB,SYT17,SLC5A7,PARK7,GRM4,NAPB,PNKD,SNAP25,VAMP1,SYT2,CRH,DRD2,CADPS,GDNF,CPLX1,TRH,BACE1,VAMP2,PRAF2* |
| Neurotransmitter secretion | *DTNBP1,SYT17,SLC5A7,GRM4,NAPB,PNKD,SNAP25,VAMP1,SYT2,DRD2,CADPS,CPLX1,BACE1,VAMP2* |
| Synaptic vesicle cycle | *DTNBP1,SNCB,AMPH,SYNDIG1,SYT17,PACSIN1,NAPB,SNAP25,VAMP1,SYT2,DRD2,CADPS,CPLX1,BACE1,VAMP2* |
| Synaptic vesicle exocytosis | *DTNBP1,SYT17,NAPB,SNAP25,VAMP1,SYT2,DRD2,CADPS,CPLX1,BACE1,VAMP2* |
| Vesicle-mediated transport in synapse | *DTNBP1,SNCB,AMPH,SYNDIG1,SYT17,PACSIN1,NAPB,SNAP25,VAMP1,SYT2,DRD2,CADPS,CPLX1,BACE1,VAMP2* |

**Supplementary table S4**: Comparison of DEGs with transcriptomic profile of the published *Setd1a*$^{+/-}$ mice.

| List of genes | |
|---|---|
| Comparison with published datasets (Mukai et al, 2019) | Upregulated genes: *SLITRK4,PDZD2,POSTN,RASGEF1B,ATP6V1F,UCP2, PARVA* |
| | Down-regulated genes: *ARID4A,FGF11,KMT2A,ZNF462* |

**Supplementary table S5**: Comparison of DEGs with SFARI datasets.

| List of genes | |
|---|---|
| Overlap with SFARI genes | *ADNP,ALDH1A3,ASTN2,ASXL3,ATP1A3,ATP2B2,CACNA2D3,CADPS,CAMK4,CDH8,CDH9,CELF6,CEP290,CHD3,CHRM3,CPEB4,CSNK2A1,DCC,DLG2,DLX6,DPYSL3,DRD2,EPC2,FABP5,FHIT,FOXG1,GLRA2,GPR37,GRIK3,GRIN2A,HLADPB1,IL1RAPL1,INPP1,KCNB1,KCND2,KCTD13,KDM5B,KIRREL3,KMT2A,LAMB1,LRRC4C,MACROD2,MAOB,MBD5,MEGF11,MET,MKX,MRTFB,MYO16,NEGR1,NEO1,NINL,NRXN3,NUAK1,PATJ,PCDHA10,PCDHA4,PCDHA6,PLCB1,PRICKLE2,RALGAPB,RBFOX1,RPS6KA2,SATB1,SH3RF3,SLC12A5,SLC25A27,SLC9A9,SNAP25,STAG1,SYT17,TSHZ3,TTN,UNC5D,VAMP2,ZMYND8,ZNF462,ZNF827* |

**Supplementary table S6:** Cell type specific electrophysiological properties of neurons in glutamatergic cultures during development. Statistical testing compared genotypes per cell class and DIV.* = based on KS-test on cumulative data.

**Glutamatergic cultures**

| **DIV 21** | Control | *SETD1A*$^{+/-}$ | |
|---|---|---|---|
| **Passive intrinsic properties** | (n = 33) | (n = 32) | p-value |
| Resting membrane potential (mV) | -36.4 ± 1.4 | -38.0 ± 1.8 | |
| Membrane resistance (MΩ) | 827 ± 64 | 967.0 ± 52 | |
| Membrane capacitance (pF) | 44.1 ± 3.0 | 32.3 ± 1.5 | <0.001 |
| **Active intrinsic properties** | (n = 18) | (n = 20) | p-value |
| AP threshold (mV) | -30.7 ± 1.4 | -29.4 ± 1.5 | |
| Rheobase (pA) | 20.1 ± 3.0 | 16.9 ± 2.2 | |
| 1st AP amplitude | 107.3 ± 1.7 | 107.8 ± 2.4 | |
| 1st AP Half time (ms) | 1.4 ± 0.1 | 1.4 ± 0.1 | |
| Adaptation rate 8-9rd/2-3rd AP | 1.4 ± 0.1 | 1.5 ± 0.1 | |
| 1st AP AHP time (ms) | 6.4 ± 0.4 | 5.6 ± 0.4 | |
| 1st-2nd AP AHP amplitude (mV) | 0.1 ± 0.4 | 0.4 ± 0.5 | |
| **Synaptic inputs/ postsynaptic events** | (n = 27) | (n = 23) | p-value |
| sEPSC amplitude (pA) | 29.3 ± 2.6 | 26.4 ± 2.2 | 0.038 |
| sEPSC frequency (Hz) | 3.4 ± 0.5 | 3.6 ± 0.8 | |
| sEPSC synchronous inputs (n/min) | 1.1 ± 0.2 | 3.2 ± 0.5 | <0.001 |

**DIV 49**

| **Passive intrinsic properties** | (n = 17) | (n = 12) | p-value |
|---|---|---|---|
| Resting membrane potential (mV) | -45.0 ± 1.9 | -47.5 ± 2.6 | |
| Membrane resistance (MΩ) | 368.8 ± 51.4 | 367.5 ± 47.4 | |
| Membrane capacitance (pF) | 56.6 ± 3.7 | 74.8 ± 7.2 | 0.020 |
| **Active intrinsic properties** | (n = 16) | (n = 11) | p-value |
| AP threshold (mV) | -33.3 ± 1.2 | -29.1 ± 0.9 | 0.020 |
| Rheobase (pA) | 84.1 ± 13.4 | 60.9 ± 0.1 | |
| 1st AP amplitude | 103.0 ± 1.3 | 98.5 ± 2.5 | |
| 1st AP Half time (ms) | 1.0 ± 0.1 | 1.2 ± 0.1 | 0.010 |
| Adaptation rate 8-9rd/2-3rd AP | 1.7 ± 0.2 | 2.1 ± 0.4 | |
| 1st AP AHP time (ms) | 3.9 ± 0.5 | 5.2 ± 0.8 | |
| 1st-2nd AP AHP amplitude (mV) | -1.0 ± 0.6 | 0.1 ± 0.8 | |
| **Synaptic inputs/ postsynaptic events** | (n = 9) | (n = 12) | p-value |
| sEPSC amplitude (pA) | 30.4 ± 2.8 | 28.1 ± 3.0 | |
| sEPSC frequency (Hz) | 1.3 ± 0.4 | 2.3 ± 0.4 | |
| sEPSC synchronous inputs (n/min) | 0.8 ± 0.3 | 5.9 ± 1.0 | <0.001 |

**Supplementary table S7**: Upregulated DEGs involve in second messenger signaling, Related to Figure 5a.

| Term | List of genes |
|---|---|
| G protein-coupled receptor signaling pathway, coupled to cyclic nucleotide second messenger | *ADCY2,PTHLH,CHGA,GSK3A,GLP1R,GHRH,GRM4,RAMP1,SSTR1,HTR7,DRD2,ADCY8,GRIK3,GPR37,ADGRG2,GRM3* |
| Adenylate cyclase-modulating G protein-coupled receptor signaling pathway | *ADCY2,PTHLH,CHGA,GSK3A,GLP1R,GHRH,GRM4,RAMP1,DRD2,ADCY8,GRIK3,GPR37,ADGRG2,GRM3* |
| Second-messenger-mediated signaling | *TMEM38A,ADCY2,PTHLH,FKBP1A,SLC9A1,CHGA,NCALD,GSK3A,GLP1R,VEGFA,NR5A2,GHRH,RAMP1,EPHA5,CRH,GSTO1,DRD2,ADCY8,PRNP,NUDT4,ADGRG2,GRIN2A* |
